# Metabolic Reprogramming by Mutant GNAS Creates an Actionable Dependency in Intraductal Papillary Mucinous Neoplasms of the Pancreas

**DOI:** 10.1101/2024.03.13.584524

**Authors:** Yuki Makino, Kimal I Rajapakshe, Benson Chellakkan Selvanesan, Takashi Okumura, Kenjiro Date, Prasanta Dutta, Lotfi Abou El-Kacem, Akiko Sagara, Jimin Min, Marta Sans, Nathaniel Yee, Megan J Siemann, Jose Enriquez, Paytience Smith, Pratip Bhattacharya, Michael Kim, Merve Dede, Traver Hart, Anirban Maitra, Fredrik I Thege

**Author notes:** Corresponding Authors: Yuki Makino, M.D., Ph.D., Department of Translational Molecular Pathology Sheikh Ahmed Center for Pancreatic Cancer Research, The University of Texas MD Anderson Cancer Center, Houston, TX, USA, Fredrik I Thege, Ph.D., Department of Translational Molecular Pathology Sheikh Ahmed Center for Pancreatic Cancer Research, The University of Texas MD Anderson Cancer Center, Houston, TX, USA. Conflict of Interest: A.M. is listed as an inventor on a patent that has been licensed by Johns Hopkins University to Thrive Earlier Detection. A.M. serves as a consultant for Tezcat Biosciences.

## Abstract

**Objective:** Oncogenic “hotspot” mutations of *KRAS* and *GNAS* are two major driver alterations in Intraductal Papillary Mucinous Neoplasms (IPMNs), which are *bona fide* precursors to pancreatic ductal adenocarcinoma. We previously reported that pancreas-specific *Kras*^G12D^ and *Gnas*^R201C^ co-expression in p48*^Cre^*; *Kras*^LSL-G12D^; Rosa26^LSL-rtTA^; Tg (TetO-*Gnas^R201C^*) mice (“*Kras;Gnas*” mice) caused development of cystic lesions recapitulating IPMNs. Here, we aim to unveil the consequences of mutant *Gnas*^R201C^ expression on phenotype, transcriptomic profile, and genomic dependencies.

**Design:** We performed multimodal transcriptional profiling (bulk RNA sequencing, single cell RNA sequencing, and spatial transcriptomics) in the “*Kras;Gnas”* autochthonous model and tumor-derived cell lines (*Kras;Gnas* cells), where *Gnas*^R201C^ expression is inducible. A genome-wide CRISPR/*Cas*9 screen was conducted to identify potential vulnerabilities in *Kras^G12D^;Gnas^R201C^*co-expressing cells.

**Results:** Induction of *Gnas*^R201C^ – and resulting G_(s)_alpha signaling – leads to the emergence of a gene signature of gastric (pyloric type) metaplasia in pancreatic neoplastic epithelial cells. CRISPR screening identified the synthetic essentiality of glycolysis-related genes *Gpi1* and *Slc2a1* in *Kras*^G12D^;*Gnas*^R201C^ co-expressing cells. Real-time metabolic analyses in *Kras;Gnas* cells and autochthonous *Kras;Gnas* model confirmed enhanced glycolysis upon *Gnas*^R201C^ induction. Induction of *Gnas*^R201C^ made *Kras*^G12D^ expressing cells more dependent on glycolysis for their survival. Protein kinase A-dependent phosphorylation of the glycolytic intermediate enzyme PFKFB3 was a driver of increased glycolysis upon *Gnas*^R201C^ induction.

**Conclusion:** Multiple orthogonal approaches demonstrate that *Kras*^G12D^ and *Gnas*^R201C^ co-expression results in a gene signature of gastric pyloric metaplasia and glycolytic dependency during IPMN pathogenesis. The observed metabolic reprogramming may provide a potential target for therapeutics and interception of IPMNs.

**SUMMARY:** *What is already known on this topic:*

- Activating “hotspot” mutations of *KRAS* and *GNAS* are found in a majority of Intraductal Papillary Mucinous Neoplasms (IPMNs).
- Expression of mutant *KRAS* and *GNAS* drives development of IPMN-like cystic lesions in the murine pancreas that eventually progress to pancreatic ductal adenocarcinoma (PDAC).

*What this study adds:*

- Mutant *GNAS* and the resulting aberrant G_(s)_alpha signaling drives a transcriptional signature of gastric (pyloric type) metaplasia in IPMNs with mucin production.
- Aberrant G_(s)_alpha signaling enhances glycolysis via protein kinase A-dependent phosphorylation of the glycolytic enzyme PFKFB3.
- Enhanced glycolysis in *KRAS;GNAS*-mutated IPMN cells is validated via multiple orthogonal approaches *in vitro* and *in vivo* and represents an actionable metabolic vulnerability.

*How this study might affect research, practice or policy:*

- The present study provides mechanistic insight into how aberrant G_(s)_alpha signaling alters the biology of *Kras*-mutant pancreatic epithelial neoplasia through metaplastic and metabolic reprogramming.
- Targeting glycolysis in IPMNs may represent both a therapeutic avenue as well as an opportunity for intercepting progression to invasive cancer.

## Introduction

The multistep progression of pancreatic ductal adenocarcinoma (PDAC) occurs via one of two *bona fide* precursor pathways, either via microscopic pancreatic intraepithelial neoplasias (PanINs) or through macroscopic cystic lesions, of which Intraductal Papillary Mucinous Neoplasms (IPMNs) are the most common [1]. The progression from IPMN to cancer accounts for approximately 10-15% of PDAC cases annually, providing an opportunity for early detection and cancer interception in this lethal disease [2]. Activating point mutations in *KRAS* are the most frequent somatic alteration in IPMNs, observed in greater than 80% of cases, followed by point mutations of *GNAS* codon 201, which is present in approximately two-thirds of IPMNs [3, 4]. Overall, approximately half of IPMNs harbor co-mutations of *KRAS* and *GNAS*, with progression to invasive carcinoma associated with additional alterations, such as *TP53*, *SMAD4* and *PIK3CA* mutations. *GNAS* encodes for the alpha subunit of a stimulatory G-protein (G_(s)_alpha protein) and the codon 201 “hotspot” mutation leads to constitutive activation due to impaired GTPase activity [5]. Activated G_(s)_alpha induces adenylyl cyclase to produce cyclic AMP (cAMP), which in turn stimulates two intracellular cAMP effectors, Protein Kinase A (PKA) and Exchange Protein directly Activated by cAMP (EPAC), impacting diverse biological functions [6, 7].

In our previous study, we demonstrated that co-expression of *Kras*^G12D^ and *Gnas*^R201C^ (p48Cre; Kras^LSL-G12D^; Rosa26^LSL-rtTA^; Tg (TetO-Gnas^R201C^), henceforth referred to as “*Kras;Gnas*” mice), caused development of IPMN-like cystic lesions in the pancreas, culminating in adenocarcinoma [8]. Thus, the “*Kras;Gnas*” autochthonous model both genocopies the most frequent combination of genetic alterations in human IPMNs, and phenocopies the observed multistep progression of IPMNs to invasive cancer. However, while this foundational study of the *Kras;Gnas* mice characterized the autochthonous model, it did not comprehensively elucidate the significance of oncogenic G_(s)_alpha signaling on tumor biology. To this end, we established murine PDAC cell lines from the *Kras;Gnas* mice (*Kras;Gnas* cells), in which *Kras*^G12D^ is constitutively expressed, while *Gnas*^R201C^ can be induced upon addition of doxycycline *in vitro*.

This provided a facile isogenic system to study the specific impact of aberrant G_(s)_alpha signaling on a mutant Ras background. Multimodal transcriptional profiling of *Kras;Gnas* cell lines and autochthonous *Kras;Gnas* mice revealed the emergence of a gene signature of gastric (pyloric type) metaplasia, defined as the upregulation of Spasmolytic Polypeptide Expressing Metaplasia (SPEM) markers (*Tff2*, *Aqp5*, *Gkn3*, etc.) with pit cell marker expression (*Tff1*, *Gkn1*, *Gkn2*, *Muc5ac*, etc.) [9, 10], upon *Gnas*^R201C^ induction. A genome wide CRISPR/*Cas9* screen conducted in *Kras;Gnas* cells identified the glycolysis-related genes *Slc2a1* and *Gpi1* as a synthetic essentiality in the setting of *Gnas*^R201C^ expression. Multiple orthogonal modalities (transcriptional profiling, Seahorse metabolic analysis and ^13^C-pyruvate hyperpolarized MR spectrometry) all confirmed enhanced glycolysis upon *Gnas*^R201C^ induction. Mechanistically, *Gnas*^R201C^ induction leads to PKA-dependent phosphorylation and consequent activation of 6-phosphofructo-2-kinase/fructose-2,6-biphosphatase 3 (p-PFKFB3), a key regulator of glycolysis [11], thus rendering the *Kras;Gnas* cells highly dependent on glycolysis. Our study identifies the role of aberrant G_(s)_alpha signaling in mucus production and metabolic reprogramming in IPMNs, which could form the basis for therapeutic and interception strategies in these PDAC precursor lesions.

## Materials and Methods

### Autochthonous Mice

The autochthonous model of IPMN, which results from pancreas-specific co-expression of mutant *Kras*^G12D^ and *Gnas*^R201C^ (*Kras;Gnas* mice) has been described [8]. Briefly, we generated *p48^Cre^*; *Kras^LSL-G12D^*; *Rosa26^LSL-rtTA^*, Tet operon (TetO)-*Gnas^R201C^* mice. In these mice, mutant *Kras* is constitutively expressed within the *p48 (Ptf1a)* positive domain, comprised of the pancreatic epithelium. To induce *Gnas*^R201C^ expression, mice were fed with 0.0060 % doxycycline diet from 8 weeks of age, which results in generation of IPMN-like cystic lesions. All animal experiments were performed in accordance with the MD Anderson Institutional Care and Use of Animals Committee (IACUC)-approved protocols.

### Establishment of cell lines from Kras;Gnas mice

We generated 4 independent tumor-derived cell lines (LGKC-1, 2, 3, and 4 cells) from *Kras;Gnas* mice for all the experiments (*Kras;Gnas* cells). Cells were established from the tumor tissues in an 8-month-old male mouse fed with normal diet (LGKC-1 cells), a 10-month-old female mouse fed with normal diet (LGKC-2 cells), a 5-month-old male mouse fed with doxycycline diet (LGKC-3 cells), and a 5-month-old female mouse fed with normal diet (LGKC-4 cells). Tissues were minced, resuspended in Roswell Park Memorial Institute (RPMI) (Corning, Corning, NY, USA) supplemented with 10 % fetal bovine serum (FBS) and 1% penicillin and streptomycin, and plated onto collagen coated dishes to expand the tumor cells [8]. Established cell lines were cultured at 37°C with 5% CO2 in RPMI supplemented with 10 % FBS and 1% penicillin and streptomycin unless otherwise indicated. All *Kras;Gnas* cell lines express mutant *Kras*^G12D^ constitutively. To induce *Gnas*^R201C^ co-expression, cells were treated with doxycycline (typically 100ng/mL), thus creating a facile isogenic system to interrogate the specific impact of G_(s)_alpha signaling on a mutant Ras background.

## Results

### Additional Methods are included as supplementary data

#### Induction of Gnas^R201C^ expression drives transcriptional reprogramming of IPMN cells with gene signatures of gastric (pyloric type) metaplasia

We established cell lines (LGKC-1, 2, 3, and 4) from PDAC arising in four independent *Kras;Gnas* mice fed with either doxycycline diet or normal diet (**Figure 1A**). To investigate transcriptional reprogramming in *Gnas*^R201C^-expressing *Kras;Gnas* cells, we performed bulk RNA sequencing in the presence (*Kras* ON, *Gnas* ON) or absence (*Kras* ON, *Gnas* OFF) of doxycycline. Approximately 90% of Gnas reads in doxycycline-treated cells harbored the R201C point mutation, whereas vehicle-treated samples had <1% mutated reads, indicating conditional induction of *Gnas*^R201C^ expression and minimal leakiness in the absence of doxycycline (**Figure 1B**). Pairwise differential expression analysis revealed 627 differentially expressed genes (p<0.05, minimum 2-fold difference) in *Gnas*^R201C^-expressing cells, of which 474 (76%) were over-expressed (**Figure 1C-D**). Gene Set Enrichment Analysis (GSEA) identified G protein-coupled receptor (GPCR) signaling, Metabolism, Coagulation/Complement, Cytoskeleton organization, and IFN/Immune response as positively enriched gene set categories (**Figure 1E and Supplementary** Figure 1). The enrichment of GPCR signaling gene sets indicates activation of downstream signaling of mutant G_(s)_alpha, consistent with *Gnas*^R201C^ induction (**Supplementary** Figure 1). Notably, transcripts of prototypal markers for SPEM (*Aqp5, Gkn3*) and gastric pit cells (*Gkn1, Gkn2, Tff1, Mucl3*), which are characteristic of gastric type IPMN [12, 13, 14], were significantly upregulated in *Gnas*^R201C^-expressing cells together with apomucins (*Muc1, Muc5b*) (**Figure 1F**). GSEA also revealed that gene signatures of gastric pit cells were enriched upon *Gnas*^R201C^ induction (**Figure 1G**). These findings indicated the emergence of a gene signature of gastric (pyloric type) metaplasia.

**Figure 1.**
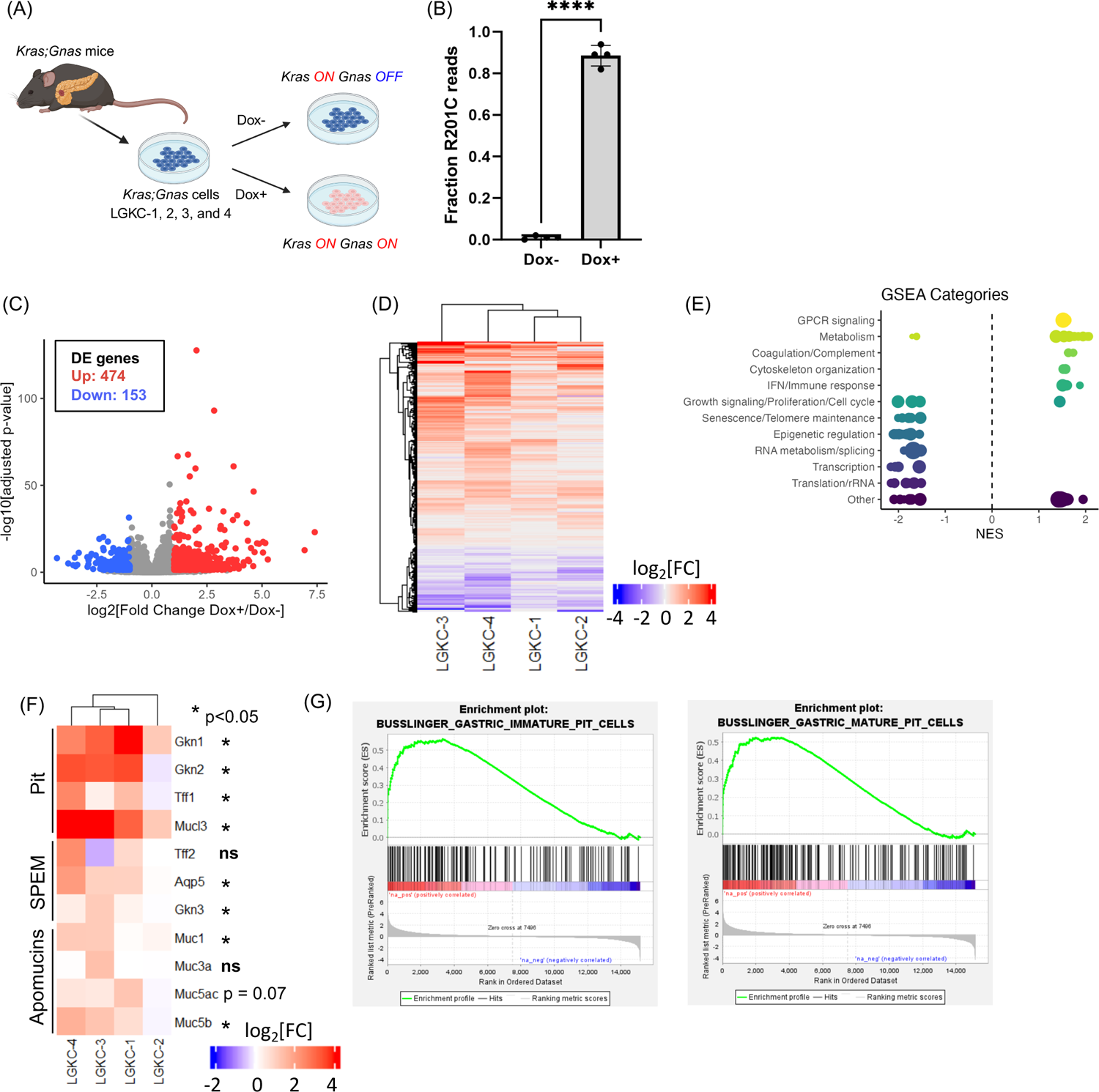
RNA-sequencing (RNA-seq) reveals transcriptional reprogramming and gene signature of gastric (pyloric type) metaplasia in *Kras;Gnas* cells with aberrant G_(s)_alpha signaling. (A) Establishment of 2D cell lines (*Kras;Gnas* cells; LGKC-1, 2, 3, and 4) from the pancreas of *Kras;Gnas* model and *in vitro* induction of *GNAS*^R201C^-expression by doxycycline (Dox). Created with BioRender. (B-G) *Kras;Gnas* cells was incubated with or without 1 ug/mL doxycycline for 24 hours before RNA collection for RNA-seq. (B) Fraction of Gnas reads harboring the R201C mutation in doxycycline-treated and untreated cells. (B) Volcano plot of RNA-seq. Significantly upregulated or downregulated genes upon doxycycline treatment were shown in red or blue, respectively. Paired analysis comparing doxycycline-positive over negative samples for four cell lines. DE; differentially expressed (C) Heatmap of differentially expressed genes upon doxycycline treatment. (D) Categories of significant genes sets enriched in doxycycline-treated cells generated through gene set enrichment analysis (GSEA). (E) Heatmap of transcripts indicative of gastric (pyloric type) metaplasia, and apomucins in doxycycline-treated cells versus untreated cells. The pyloric type metaplasia signature includes transcripts indicative of both gastric pit and Spasmolytic Polypeptide Expressing Metaplasia (SPEM). (F) GSEA showing the enrichment of gastric pit cell gene signatures in doxycycline-treated cells over untreated cells.

To further analyze the transcriptional alterations in pancreatic epithelial cells *in vivo,* we performed single cell RNA sequencing (scRNA-seq) in *Kras;Gnas* mice fed with doxycycline (*Kras ON, Gnas ON*) *versus* normal diet (*Kras ON, Gnas OFF*) for 10 weeks. In both cohorts, we identified a heterogenous pancreatic epitthelial compartment, consisting of seven distinct clusters (**Figure 2A**). Of these, three epithelial clusters were observed in both cohorts and identified as ‘acinar’, ‘duct-like’, and ‘ADM-like’, characterized by high expression of acinar-specific genes (*Cpa1, Cela2a*), canonical ductal transcription factors (*Hnf1b, Onecut1, Sox9*), and co-expression of acinar and ductal markers, respectively (**Figure 2B-C**). On the contrary, four epithelial clusters were almost exclusively present in the mice fed with doxycycline diet, suggesting these populations were driven by aberrant G_(s)_alpha signaling (**Figure 2B**). Among the four, three clusters were “duct-like” in origin based on *Krt19* and *Car2* co-expression (**Figure 2C**). These three “duct-like” clusters were also characterized by elevation of well described markers of metaplasia (*Onecut2, Foxq1*) [12, 15, 16]. The fourth was identified as a minor tuft cell cluster based on co-expression of *Dclk1* and *Pou2f3* [15]. Importantly, these three metaplastic “duct-like” clusters were distinct from the ADM-like cluster, suggesting they were specifically accentuated by aberrant G_(s)_alpha signaling. Of the three metaplastic “duct-like” clusters, one additionally displayed elevated gastric pit cell markers, and is referred to as “metaplastic pit-like”. The other had elevated markers of proliferation (*Mki67, Pcna*), which we refer to as “metaplastic duct-like proliferating”. SPEM markers were most abundant in the “metaplastic duct-like” cluster. The three metaplastic “duct-like” clusters also displayed relative overexpression of transcripts encoding for apomucins (*Muc6, Muc5ac, Muc4*) that are associated with gastric type IPMNs (**Figure 2D**) [17]. These findings suggested the induction of *Gnas*^R201C^ expression provoked gastric (pyloric type) metaplasia and associated mucin production.

**Figure 2.**
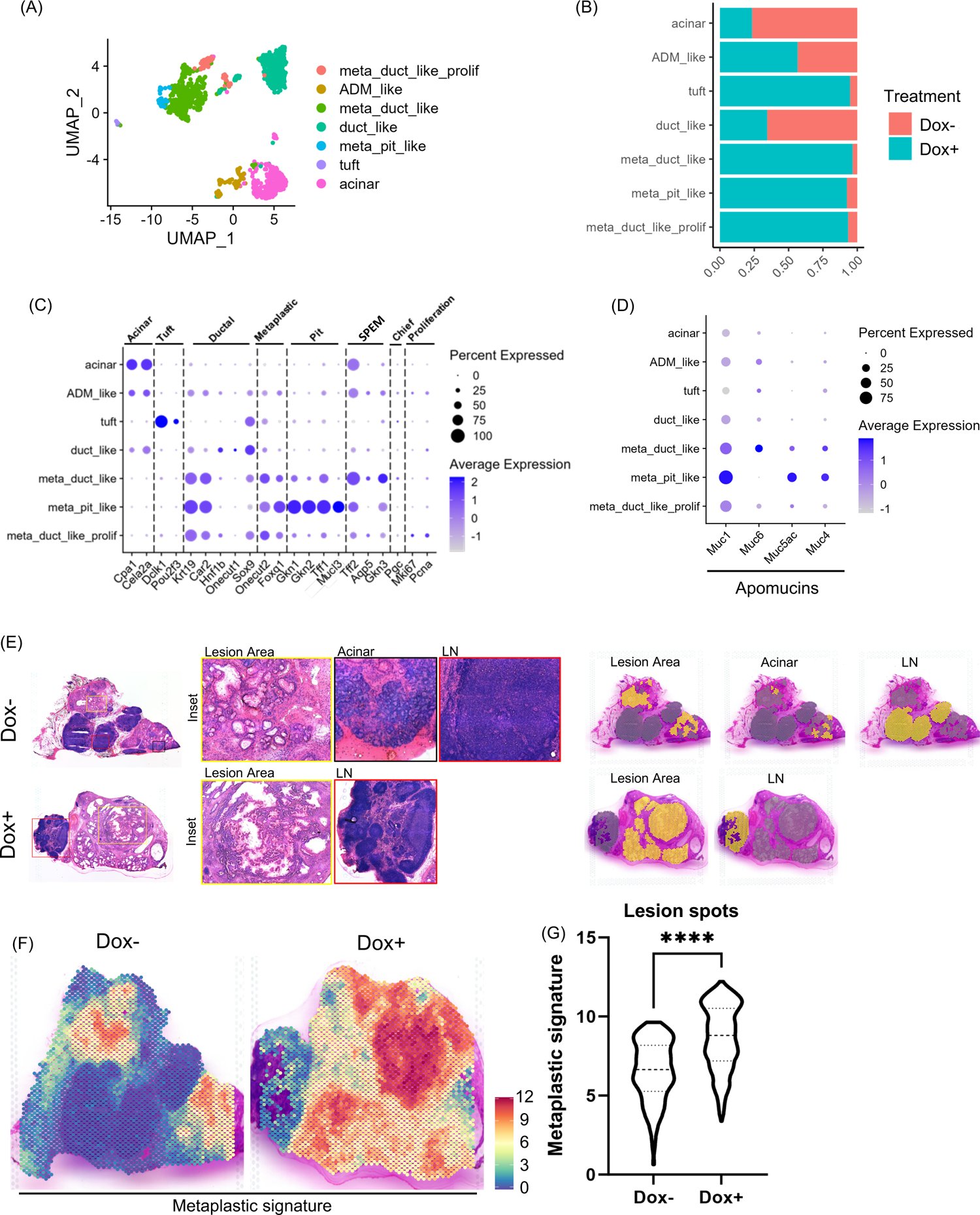
Single-cell RNA-sequencing (scRNA-seq) and spatial transcriptomics (ST) identifies heterogeneous metaplastic duct-like populations with gastric (pyloric type) metaplasia in *Kras;Gnas* mice with aberrant G_(s)_alpha signaling. (A-D) scRNA-seq was performed in the pancreas in *Kras;Gnas* mice fed with doxycycline (*Kras ON, Gnas ON*) *versus* normal diet (*Kras ON, Gnas OFF*) for 10 weeks (N = 2 for each group). (A) Uniform Manifold Approximation and Projection (UMAP) plot of seven distinct epithelial cell clusters. (B) Proportion of cells in epithelial subclusters in doxycycline fed *versus* normal diet fed *Kras;Gnas* mice. (C) Dot plots showing the expression of representative annotation markers in each epithelial cell cluster. ‘Metaplastic duct-like’, ‘metaplastic pit-like’, and ‘metaplastic duct-like proliferating’ clusters showed upregulation of ductal and metaplastic markers indicative of duct-like origin with metaplastic characteristics. ‘Metaplastic duct-like’ and ‘metaplastic pit-like’ clusters represent gastric (pyloric) metaplasia as evidenced by the upregulation of SPEM and pit cell markers, respectively. (D) Expression of transcripts for cellular apomucins in epithelial cell clusters demonstrates enrichment within metaplastic clusters. (E-G) ST of pancreatic tissues in *Kras;Gnas* mice fed with normal diet or doxycycline diet for 25 weeks (N = 1 per group). Data sets from the doxycycline-fed mouse was obtained from Reference 12 (Sans, et al. Cancer Discovery 2023). (E) Spatial mapping of the “Lesion”, “Acinar”, and “Lymph Node” (LN) areas in the pancreatic tissues in *Kras;Gnas* mice. (F) Spatial analysis of the enrichment of metaplastic gene signature in *Kras;Gnas* mice fed with doxycycline *versus* normal diet. The Metaplastic gene signature consists of 13 genes associated with gastric (pyloric type) metaplasia analyzed in our bulk and single-cell RNA-seq data sets (*Gkn1, Gkn2, Gkn3, Tff1, Tff2, Aqp5, Mucl3, Muc1, Muc3a, Muc5ac, Muc5b, Onecut2, Foxq1*). (G) Upregulation of a metaplastic gene signature in lesion spots in the doxycycline-fed *Kras;Gnas* mouse over normal diet-fed *Kras;Gnas* mouse. ****p<0.0001.

To validate the histological relevance of the gene signatures, we next analyzed spatial transcriptomics (ST) in the *Kras;Gnas* mice fed with doxycycline diet or normal diet, using the data from our recently published study [12]. We manually annotated areas in the tissue sections as “Lesion”, “Acinar”, and “Lymph Node (LN)” based on morphological examination (**Figure 2E**). We calculated a “Metaplastic signature” score for each spot based on the expression of 13 genes associated with gastric (pyloric type) metaplasia in our bulk and single-cell RNA-seq data sets (*Gkn1, Gkn2, Gkn3, Tff1, Tff2, Aqp5, Mucl3, Muc1, Muc3a, Muc5ac, Muc5b, Onecut2, Foxq1*). This score was significantly higher in the lesion spots from the doxycycline-treated mouse and specifically correlated with areas with IPMN-like histology (**Figure 2F-G**).

In summary, multimodal profiling, including bulk RNA-seq on cell lines and scRNA-seq and ST in the autochthonous *Kras;Gnas* model identified that induction of aberrant G_(s)_alpha signaling leads to gastric pyloric metaplasia, resulting in a heterogenous population of metaplastic duct-like cells with characteristics of IPMN.

#### Functional genomics screen implicates glycolysis as a dependency in Gnas^R201C^-expressing Kras;Gnas cells

To interrogate genetic dependencies in IPMN cells in an unbiased fashion, we performed genome-wide CRISPR/*Cas*9 loss-of-function screening to identify genes required for survival of isogenic *Kras;Gnas* cells with aberrant G_(s)_alpha signaling. As shown in **Figure 3A**, LGKC-1 and LGKC-3 cells were transduced with a genome-wide lentiviral gRNA library and cultured with or without doxycycline treatment for 9 days. We found that gRNAs for 3 and 50 genes were significantly depleted on *Gnas*^R201C^ induction in LGKC-1 and LGKC-3 cells, respectively (**Figure 3B-C**). gRNAs targeting 3 genes (*Gpi1, Slc2a1*, and *Zfp120)* were commonly depleted in doxycycline-treated samples of both cell lines. Interestingly, *Slc2a1* and *Gpi1* encode for glucose transporter type 1 (GLUT1) and glucose-6-phosphate isomerase 1 (GPI1), respectively. GLUT1 is a glucose transporter, while GPI1 is an enzyme that catalyzes the conversion of glucose-6-phosphate to fructose-6-phosphate [18, 19], both of which are essential genes in the glycolytic pathway. These results indicated a potential dependence on glycolysis in *Kras;Gnas* cells upon induction of aberrant G_(s)_alpha signaling.

**Figure 3.**
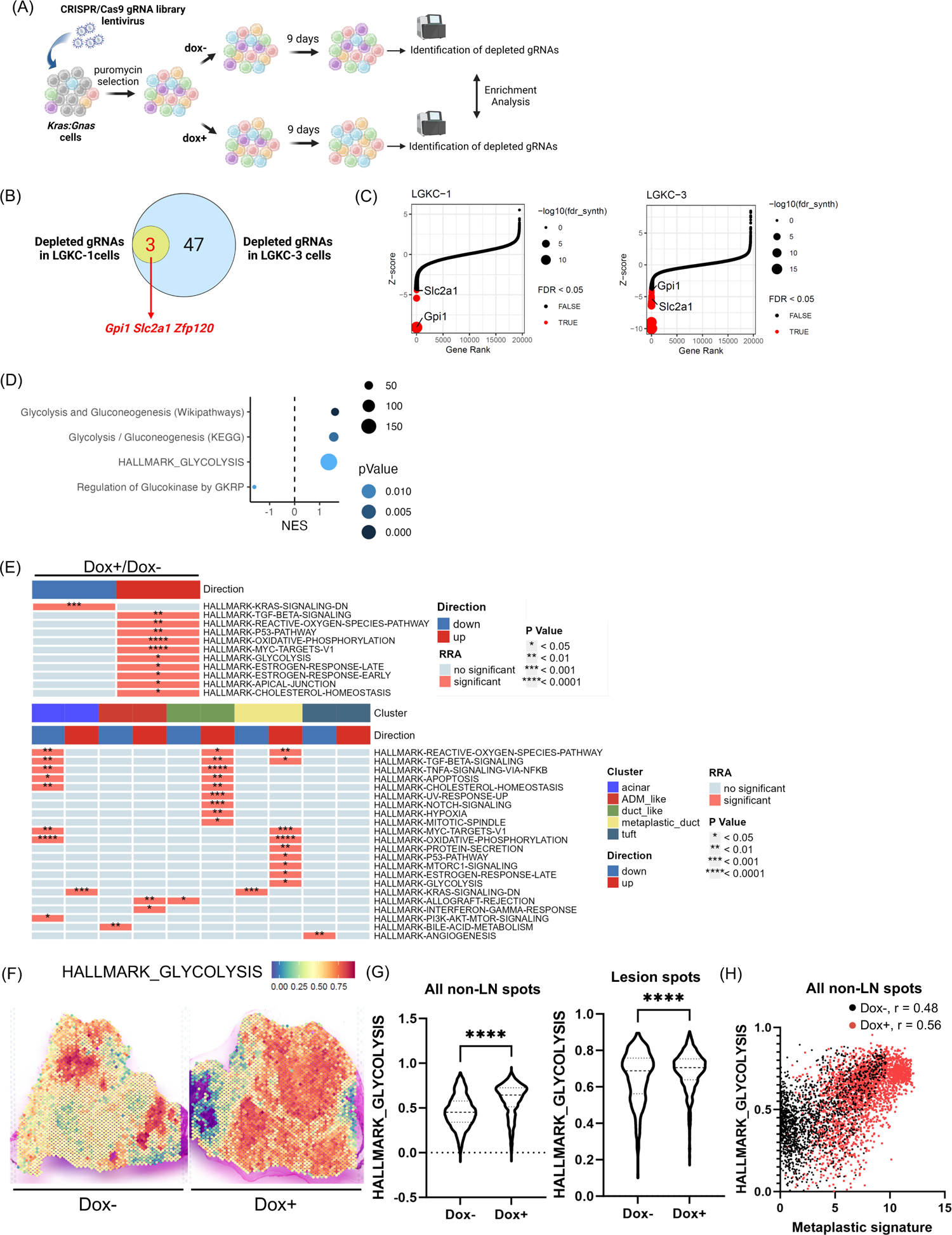
CRISPR/Cas9 loss-of-function screening identifies glycolysis as an actionable vulnerability in *Kras;Gnas* cells with aberrant G_(s)_alpha signaling. (A) Schematic illustration of genome-wide CRISPR/*Cas9* knockout screening of *Kras;Gnas* cells. Created with BioRender. (B) Venn diagram of shared depleted genes in the LGKC-1 and LGKC-3 cell line CRISPR screen. Created with BioRender. (C) CRISPR drop-out screening Z-scores of the relative abundance of guides for each gene in LGKC-1 and LGKC-3 cells cultured with vs without doxycycline. Genes with FDR <0.05 were highlighted in red in cells. (D) Glycolysis-related gene sets significantly enriched in bulk RNA-seq of doxycycline-treated *Kras;Gnas* cells *versus* untreated cell lines. (E) scGSEA of pancreatic tissues in *Kras;Gnas* mice fed with normal or doxycycline diet. scRNA-seq data shown in Figure 2 was used for the analysis. scGSEA on HALLMARK gene sets in all epithelial cells as a function of doxycycline treatment (upper) or as a function of cluster identity (lower) in *Kras;Gnas* mice. The ‘metaplastic duct-like’, ‘metaplastic pit-like’ and ‘metaplastic duct-like proliferating’ clusters shown in Figure 2 were combined into a single ‘metaplastic_duct’ cluster. (F-H) ST of the pancreatic tissues in *Kras;Gnas* mice shown in Figure 2. (F) Spatial analysis of the enrichment of HALLMARK GLYCOLYSIS gene signature in *Kras;Gnas* mice fed with doxycycline *versus* normal diet. (G) Upregulation of HALLMARK GLYCOLYSIS gene signature in all non-LN spots and lesion spots in doxycycline-fed *Kras;Gnas* mouse over normal diet-fed *Kras;Gnas* mouse. (H) Positive correlation between metaplastic gene signature and HALLMARK GLYCOLYSIS gene signature in all non-LN spots. *p<0.05, **p<0.01, ***p<0.001, ****p<0.0001.

Interrogation of bulk RNA-seq data in *Kras;Gnas* cells showed that glycolysis-related gene signatures were significantly enriched by the induction of *Gnas^R201C^* expression, including HALLMARK_GLYCOLYSIS (**Figure 3D**). To determine if these results are reproduced *in vivo*, we performed rank-based gene set enrichment analysis on our scRNA-seq data set. This identified multiple HALLMARK gene sets significantly enriched in the epithelial clusters in doxycycline-treated mice, including HALLMARK_GLYCOLYSIS (**Figure 3E, upper**). We next combined all the metaplastic cell clusters which were almost exclusively found in the doxycycline-treated mice as shown above, into one “metaplastic_duct” cluster and repeated the enrichment analysis. This revealed that the HALLMARK_GLYCOLYSIS signature was enriched only in the metaplastic cluster in the scRNA-seq dataset (**Figure 3E, lower**). In the existing ST dataset, the HALLMARK-GLYCOLYSIS signature was significantly higher in epithelial spots (excluding lymph nodes), and lesion spots in the doxycycline-treated mouse pancreas (**Figure 3F-G**). We also found a significant correlation between the Metaplastic signature and the HALLMARK_GLYCOLYSIS signature in these samples (**Figure 3H**), suggesting a putative link between these processes.

To validate these observations in human IPMNs, we analyzed the ST data from our recently published series [12]. Since gastric pit and SPEM markers are characteristics for low grade (gastric) IPMNs, we used the data of 7 low grade (gastric) IPMNs from the data set. As detailed in the previous report, three different regions - “Epilesional” (spots overlapping with the neoplastic epithelium), “Juxtalesional” (defined as two layers of spots from the neoplastic epithelium corresponding to the immediately adjacent microenvironment), and “Perilesional” (spots distal from these regions) were extracted for the analysis (**Figure 4A**). Not surprisingly, Epilesional spots showed the highest expression of transcripts corresponding to metaplasia and gastric cell markers, and apomucins (**Figure 4B**). The Metaplastic gene signature was the highest in the Epilesional spots (**Figure 4C**). We found that Epilesional spots showed the highest HALLMARK GLYCOLYSIS scores, followed by a gradual decrease in Juxtalesional and Perilesional spots (**Figure 4D**). The Metaplastic gene signature showed significant positive correlation with the HALLMARK GLYCOLYSIS signature (**Figure 4E**).

**Figure 4.**
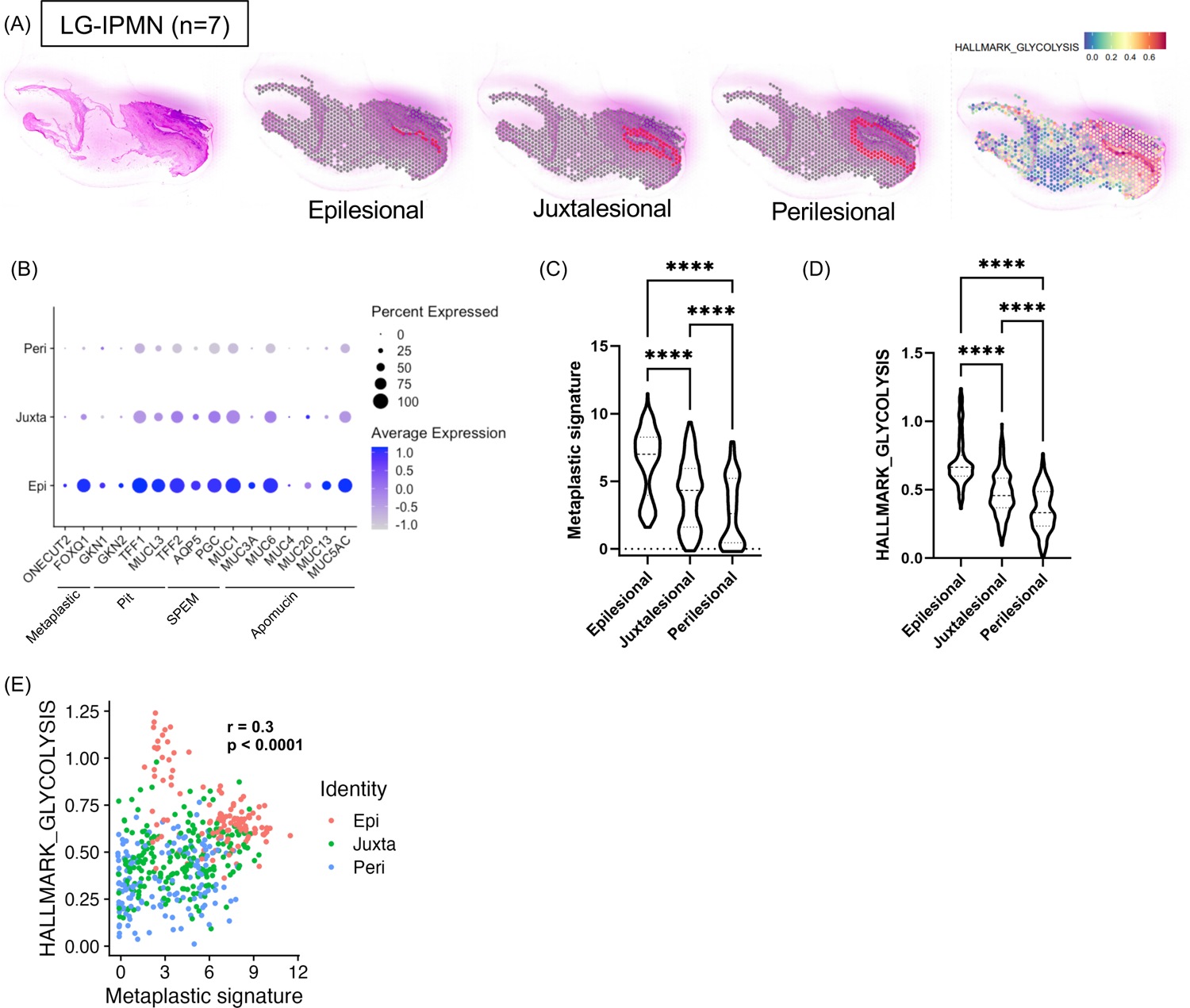
Spatial transcriptomics of human IPMNs validates glycolysis signature within neoplastic epithelium concomitant with metaplastic signature. Data set of 7 human low-grade IPMN samples were obtained from Reference 12 (Sans, et al. Cancer Discovery 2023). (A) Mapping of Epilesional, Juxtalesional, and Perilesional areas, and HALLMARK-GLYCOLYSIS genes signature in a representative low-grade IPMN sample. (B) Dot plot of the expression of metaplastic and IPMN markers as a function of tissue region. (C) Enrichment of the Metaplastic gene signature in Epilesional spots relative to Juxtalesional and Perilesional spots. (D) Enrichment of the HALLMARK GLYCOLYSIS gene signature in Epilesional spots relative to Juxtalesional and Perilesional spots. (E) Scatter plot showing the correlation between the Metaplastic and HALLMARK GLYCOLYSIS gene signature scores.

In summary, CRISPR/*Cas9* screening and cross-species transcriptomic analyses cumulatively suggest that glycolysis is potentially essential for the survival of IPMN-derived tumor cells, that induction of *Gnas*^R201C^ drives a glycolytic signature *in vitro* and *in vivo*, and that this signature is elevated in human IPMNs. We next sought to determine if *Gnas*^R201C^ expression drives an increase in glycolytic flux.

#### Induction of Gnas^R201C^ expression enhances glycolysis on real time metabolic assessment in vitro and in vivo

We next performed functional metabolic analysis to investigate the relationship between *Gnas*^R201C^ expression and glycolytic flux. Induction of *Gnas*^R201C^ expression significantly increased glucose uptake and lactate secretion in *Kras;Gnas* cells *in vitro* (**Figure 5A-B** and **Supplementary** Figure 2). We also found that the Proton Efflux Rate (PER) was elevated on *Gnas*^R201C^ induction, consistent with increased basal glycolysis (**Figure 5C**). To directly assess the glycolytic flux, we performed real-time Hyperpolarized Magnetic Resonance Spectroscopy (HP-MRS) in which the conversion of ^13^C-labeled pyruvate into lactate is measured as a lactate / pyruvate signal ratio (**Figure 5D**) [20, 21]. All four cell lines showed an increase of lactate / pyruvate signal ratio following *Gnas*^R201C^ induction, suggesting increased glycolytic flux (**Figure 5E-F**). To assess glycolytic flux *in vivo* we performed ^13^C-HP-MRS coupled with high-resolution T2-weighted proton (^1^H) MRI in *Kras;Gnas* mice. Compared with conventional diet-fed mice, doxycycline-fed mice showed elevation of the lactate / pyruvate signal ratio, consistent with elevated pancreatic glycolytic flux *in vivo* (**Figure 5G-H**). These findings provided additional lines of evidence that induction of mutant G_(s)_alpha on a mutant *Kras* background drives glycolysis in the pancreata of *Kras;Gnas* mice. Based on these findings, we subsequently analyzed the functional significance of abrogating glycolysis in IPMN models.

**Figure 5.**
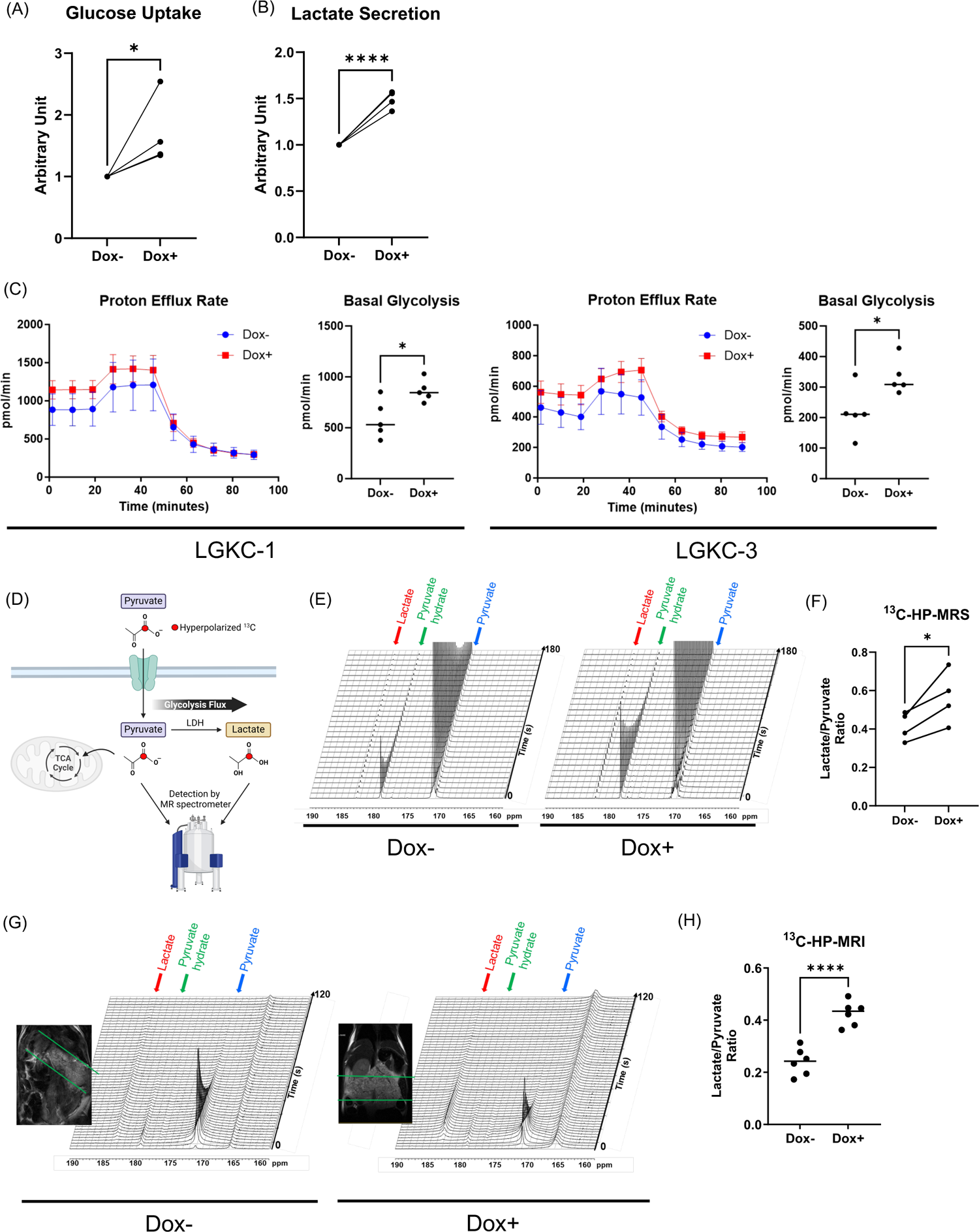
Real-time metabolic analysis identifies increased glycolytic flux following induction of mutant *GNAS in vitro* and *in vivo*. (A-B) Four different *Kras;Gnas* cells (LGKC-1, 2, 3, and 4) were independently analyzed in each assay. Cells were incubated with or without 100 ng/mL doxycycline for 24 hours before the assay. The average value of 4 replicates represented each cell line. The results of 4 replicates were shown in Supplementary Figure 2. (A) Glucose uptake from culture media into *Kras;Gnas* cells incubated with or without doxycycline. (B) Lactate secretion into culture media from *Kras;Gnas* cells incubated with or without doxycycline. (C) Real-time cell metabolic analysis with Seahorse XF Glycolytic Rate Assay to determine the effect of *GNAS*^R201C^ induction on basal glycolysis. Cells were incubated with or without 100 ng/mL doxycycline for 24 hours before the assay. Proton Efflux Rate (PER) was sequentially measured under basal conditions, after inhibition of oxidative phosphorylation by rotenone/ antimycin A (Rot/AA), and after inhibition of glycolysis by 2-DG. (D-H) Real-time cell metabolic analysis with ^13^C-pyruvate hyperpolarized magnetic resonance spectroscopy (MRS) (D) Scheme of ^13^C-pyruvate hyperpolarized MRS which detects Lactate / Pyruvate signal ratios to calculate glycolytic flux. Created with BioRender. (E) Representative sequential NMR spectra after the addition of approximately 8.7 mM hyperpolarized pyruvate in *Kras;Gnas* cells incubated with or without doxycycline for 24 hours. The spectrum was collected for every 6 seconds for 180 seconds. The labeled peaks are pyruvate, pyruvate hydrate and lactate. (F) Lactate / Pyruvate signal ratios of *Kras;Gnas* cells with or without doxycycline treatment. LGKC-1, 2, 3, and 4 cells were independently incubated with or without doxycycline for 24 hours before each assay. (G) Representative T2-weighted MRI (coronal slice) images and real-time in vivo ^13^C-magnetic resonance spectra after intravenous injection of hyperpolarized pyruvate. The sequential spectra are collected for every 2 seconds for 120 seconds from the MRI slabs on the mouse pancreas. (H) Lactate / Pyruvate signal ratios of the pancreas in *Kras;Gnas* mice on ^13^C-pyruvate Hyperpolarozed MRI. *Kras;Gnas* mice fed with normal diet or doxycycline diet for 3-15 weeks were analyzed (N = 6 for each group). *p<0.05, ****p<0.0001.

#### Glycolysis is an actionable vulnerability in Gnas^R201C^-expressing Kras;Gnas cells

To analyze the significance of glucose metabolism on cell proliferation in *Kras;Gnas* cells, we evaluated the influence of glucose deprivation. In glucose-replete medium, induction of *Gnas*^R201C^ did not impact cell proliferation (**Supplementary** Figure 3A) or colony formation (**Supplementary** Figure 3B). In contrast, induction of *Gnas*^R201C^ significantly attenuated the proliferation and colony formation of both LGKC-1 and LGKC-3 cells in glucose-free medium relative to isogenic controls (**Supplementary** Figure 3A-B). We next treated *Kras;Gnas* cells with several preclinical inhibitors of glycolysis, including WZB-117, PFK-15, and 2-Deoxy-D-glucose (2-DG). Both cell lines displayed increased sensitivity to these inhibitors upon *Gnas*^R201C^ induction as indicated by decreased half maximal inhibitory concentration (IC_50_) values in doxycycline-treated cells (**Supplementary** Figure 3C). These results indicate that *Kras;Gnas* cells expressing *Gnas*^R201C^ have greater dependency on glucose and glycolysis for their proliferation and survival, in comparison to isogenic controls.

To further validate the glycolysis dependency using data from our CRISPR/*Cas9* screen (**Figure 3A-C**), we generated two independent clones with deletion of endogenous *Gpi1* from two *Kras;Gnas* cell lines (**Figure 6A**). As expected, knockout of *Gpi1* resulted in drastically decreased basal glycolysis in these cells (**Figure 6B**). In *Kras;Gnas* cells with retained *Gpi1*, there was no difference in proliferation rate or colony formation on *Gnas*^R201C^ induction (**Figure 6C-D**). In contrast, in both sets of *Gpi1*-knockout clones, cell proliferation and colony formation were significantly reduced on *Gnas*^R201C^ induction (**Figure 6C-D**). These *in vitro* findings were recapitulated in subcutaneous allografts in athymic mice. There was no significant difference in growth between *Kras;Gnas* tumors with retained *Gpi1* expression, irrespective of normal or doxycycline diet (**Figure 6E-G**). Meanwhile, allograft volumes were significantly reduced upon *Gnas*^R201C^ induction in *Gpi1*-knockout cells (**Figure 6E-G**). Interestingly, *Gpi1* knockout resulted in downregulation of several gastric pit cell and SPEM markers in *Gnas*^R201C^-expressing cells, suggesting a potential link between glycolysis and transcriptomic programs of pyloric metaplasia (**Figure 6H**).

**Figure 6.**
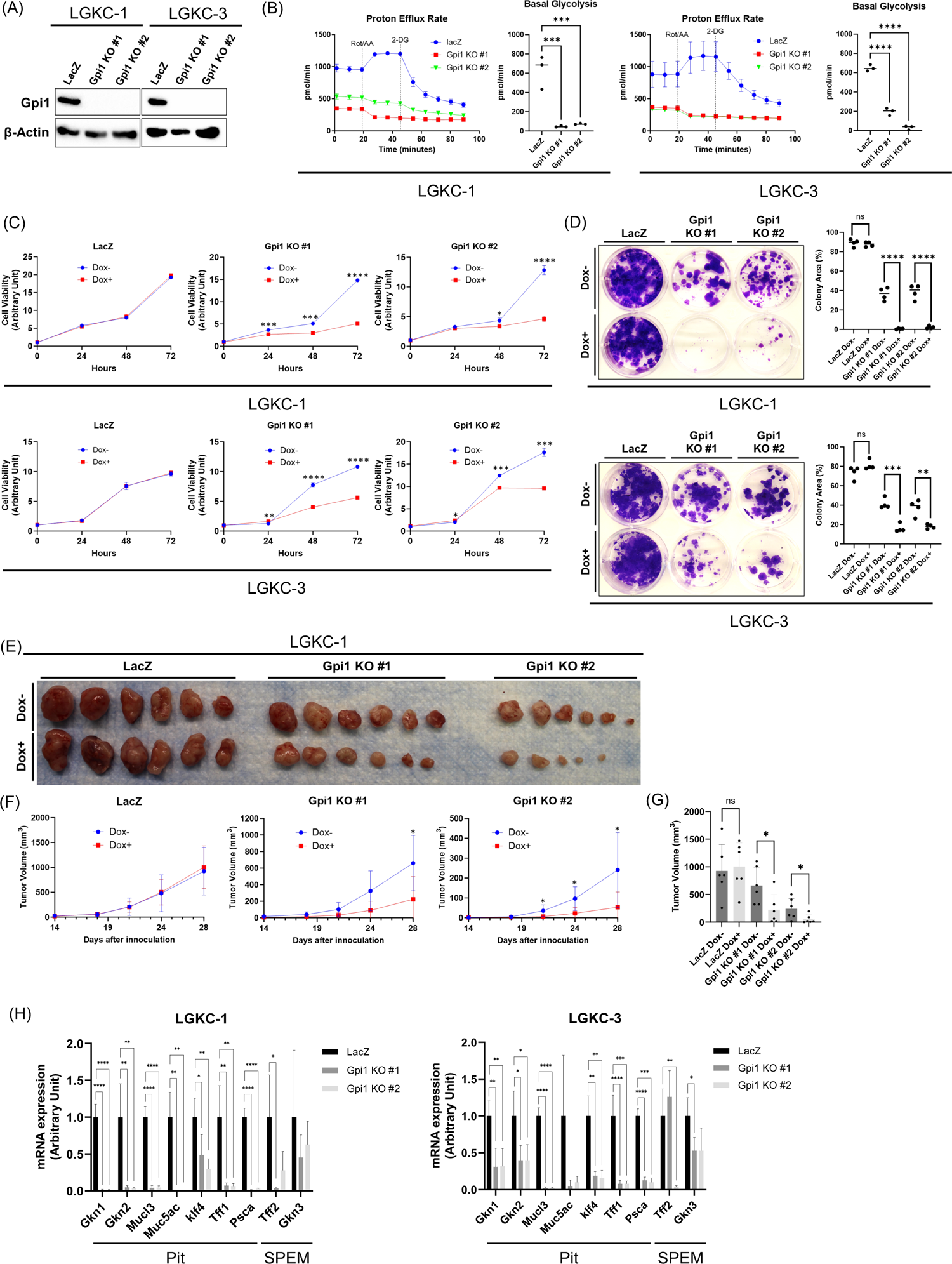
Loss of *Gpi1* abolished glycolysis and attenuated proliferation in *Gnas^R201C^*-expressing *Kras;Gnas* cells. (A) Immunoblot of CRISPR *Gpi1* KO *Kras;Gnas* cells and controls (*LacZ*). Cells were incubated with 100 ng/mL doxycycline for 24 hours before protein collection. (B) Seahorse XF Glycolytic Rate Assay. Cells treated with 100 ng/mL doxycycline for 24 hours were incubated in the medium containing glucose, glutamine, and pyruvate, followed by the injection of Rot/AA and 2-DG. PER was sequentially measured at indicated time points. (C) Cell proliferation assay. Cell viability was measured by WST-8 assay at indicated time points. Cells were incubated with or without 100 ng/mL doxycycline under glucose replete media. N = 3, technical replicates. (D) Colony formation assay. Cells were stained after 10-day incubation. Cells were incubated with or without 100 ng/mL doxycycline under glucose replete media. (E-G) Allograft injection model of *Gpi1* knockout cells. Cells were subcutaneously inoculated to the bilateral flank portion of nude mice (N = 6 tumors per group). Mice were fed with normal diet or doxycycline diet from the day of inoculation. Tumor volumes were measured every 3 or 4 days from day 14 until sacrifice on day 28. (E) Macroscopic appearance of the allograft tumors. (F) Sequential tumor volume after cell inoculation. (G) Tumor volume at the time of sacrifice (day 28). (H) Quantitative PCR analysis for gastric pit cell and spasmolytic polypeptide expressing metaplasia (SPEM) markers in *Gpi1*-knockout *Kras;Gnas* cells. Cells were incubated with 100 ng/mL doxycycline for 24 hours before RNA collection. N = 4, technical replicates. *p<0.05, **p<0.01, ***p<0.001, ****p<0.0001.

We also tested the effect of the deletion of *Slc2a1* in the setting of *Gnas*^R201C^ induction. Glut1, which is encoded by *Slc2a1*, was depleted in the knockout cells and basal glycolysis was significantly reduced (**Supplementary** Figure 4A-B). Induction of *Gnas*^R201C^ attenuated in vitro cell proliferation and colony formation of *Slc2a1*-knockout cells, compared to isogenic control lines (**Supplementary** Figure 4C-D). In the subcutaneous allograft model, doxycycline diet specifically suppressed the growth of *Slc2a1*-knockout cells (**Supplementary** Figure 4E-G). Similarly to *Gpi1* knockout, loss of *Slc2a1* decreased the expression of gastric pit cell and SPEM markers (**Supplementary** Figure 4H). Thus, our *in vitro* and *in vivo* single gene knockout studies of *Gpi1* and *Slc2a1* validated the CRISPR screen.

In summary, these findings confirm that expression of *Gnas*^R201C^ increases the requirement of glucose transport and glycolytic flux for cell proliferation in *Kras* mutant cells and that abrogation of glycolysis is a potentially actionable metabolic weakness in these cells.

#### PKA-mediated PFKFB3 phosphorylation drives enhanced glycolysis in Kras;Gnas cells on Gnas^R201C^ induction

We next investigated the mechanism of increased glycolysis by *Gnas^R201C^* induction. Induction of *Gnas^R201C^* resulted in the upregulation of phospho-PKA substrates, indicating the activation of G_(s)_alpha/cAMP/PKA signaling (**Figure 7A, left**). Expression of multiple components of the glycolytic pathway, including HK1, HK2, LDHA, GAPDH, PFKP, PKM2, and pyruvate dehydrogenase was not altered in *Kras;Gnas* cells on *Gnas^R201C^* induction (**Figure 7A, middle**). On the contrary, phosphorylation of PFKFB3, one of the phosphofructokinase (PFK) subunits that mediate the phosphorylation of fructose 6-phosphate to fructose-1,6-bisphosphate, was increased in both *Kras;Gnas* cell lines (**Figure 7A, right**). Treatment with forskolin, an agonist of adenylate cyclase, recapitulated p-PFKFB3 upregulation, together with the increase of phospho-PKA substrates, in *Kras;Gnas* cells (**Figure 7B**). Exposure to a preclinical grade PKA inhibitor H-89 [22] resulted in the attenuation of the PFKFB3 phosphorylation observed on *Gnas^R201C^*induction (**Figure 7C**), suggesting that *Gnas^R201C^*-mediated PFKFB3 phosphorylation is PKA dependent. We subsequently analyzed the impact of modulating PFKFB3 activity on glycolysis in *Gnas^R201C^*-expressing *Kras;Gnas* cells. The increase in glucose uptake and lactate secretion on *Gnas^R201C^* induction (**Figure 5A-B**) was significantly reduced by exposure to PFK-15, a preclinical grade inhibitor of PFKFB3 [11] (**Figure 7D-E**). The upregulation of basal glycolysis by *Gnas^R201C^* induction was abolished by PFK-15 (**Figure 7F**). The basal glycolytic rate in *Gnas^R201C^*-expressing cells was more sensitive to PFKFB3 inhibition by PFK-15, as indicated by lower IC_50_ values in both cell lines (**Figure 7G**). Immunohistochemistry for phosphorylated PFKFB3 in human IPMN samples (7 *GNAS*-wild and *GNAS*-mutant samples, respectively) revealed significantly increased staining in *GNAS*-mutant relative to *GNAS*-wild type lesions (**Figure 7H**). Overall, these findings support that activation of the cAMP-PKA-pPFKFB3 axis is one avenue through which induction of mutant *GNAS* enhances glycolysis in IPMNs (**Figure 7I**).

**Figure 7.**
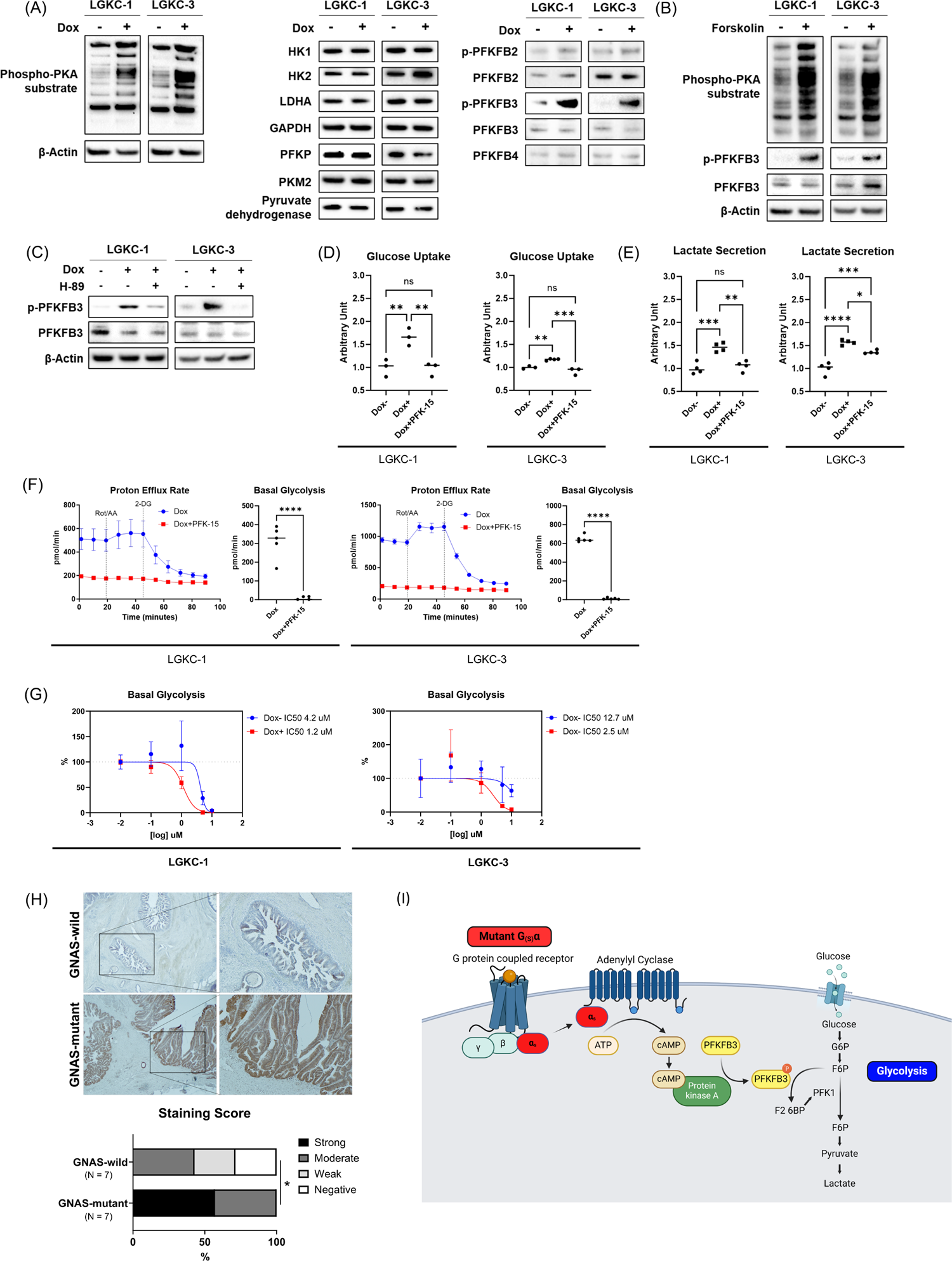
PKA-mediated activation of PFKFB3 is responsible for glycolysis enhancement induced by mutant GNAS. (A) Immunoblot of glycolysis pathway-related molecules in *Kras;Gnas* cells. Cells were incubated with or without 100 ng/mL doxycycline for 24 hours before protein collection. (B) Immunoblot in *Kras;Gnas* cells incubated with or without 10 μM forskolin for 1 hour before protein collection. (C) Immunoblot in *Kras;Gnas* cells incubated with or without 100 ng/mL doxycycline or 5 μM H-89 for 24 hours before protein collection (D-E) *Kras;Gnas* cells were incubated for 24 hours before each assay with or without 100 ng/mL doxycycline or 10 μM PFK-15. (D) Glucose uptake assay in untreated, doxycycline treated or doxycycline + PFK-15 treated *Kras;Gnas* cells. (E) Lactate secretion assay in untreated, doxycycline treated or doxycycline + PFK-15 treated *Kras;Gnas* cells. (F) Seahorse XF Glycolytic Rate Assay of *Kras;Gnas* cells. Cells were incubated with or without 10 μM PFK-15 under 100 ng/mL doxycycline for 24 hours before the assay. During the assay, cells were incubated in the medium containing glucose, glutamine, and pyruvate, followed by the injection of Rot/AA and 2-DG. PER was measured at indicated time points. (G) Dose-response curve and the half maximal inhibitory concentration (IC_50_) for basal glycolysis on PFK-15 treatment. *Kras;Gnas* cells were treated with PFK-15 at indicated concentrations with or without 100 ng/mL doxycycline treatment for 24 hours before the assay. Basal glycolysis was measured by Seahorse XF Glycolytic Rate Assay. (H) Immunohistochemistry for phospho-PFKFB3 in human IPMN with or without *GNAS* mutation (N = 7 for each group). Representative images of *GNAS*-wild and mutant cases were shown. (I) Scheme of increased glycolysis via mutant G_(s)_alpha-PKA-PFKFB3 axis. Created with BioRender. *p<0.05, **p<0.01, ***p<0.001, ****p<0.0001.

## Discussion

In the present study, multimodal transcriptome analyses of the *Kras;Gnas* IPMN model demonstrated that induction of mutant *GNAS* expression, on a mutant *KRAS* background, reprogrammed epithelial cells toward gastric (pyloric type) metaplasia. Pyloric metaplasia is a repair process that occurs in response to mucosal injury in the stomach, which involves foveolar (pit) cell hyperplasia and establishment of SPEM cells at the base of glands, following parietal cell loss [9, 10]. While pyloric metaplasia protects against ongoing mucosal injury through production of protective mucins and wound-healing agents (e.g. Tff2), it can also be an initial step towards dysplasia and tumorigenesis [9, 10]. Interestingly, analogous gastric pit and SPEM markers are also characteristic of low grade (gastric) IPMNs, which are the most common subtype of IPMNs [23, 24, 25]. Our findings demonstrate that mutant *GNAS* at least partially contributes to the gastric metaplastic phenotype characteristic of low-grade IPMNs. We found that mutant *GNAS* exacerbates pyloric metaplasia and drives the development of gastric-like IPMNs. We have previously described the expression of the transcription factor *Nkx6-2* in low-grade IPMNs, and the association between *Nkx6-2* upregulation and the gastric (pyloric like) signature [12]. Surprisingly, in our transcriptomic analyses, we did not observe upregulation of Nkx6-2 on induction of *Gnas^R201C^* (*data not shown*). suggesting that this pivotal transcription factor is induced independent of G_(s)_alpha signaling in early IPMN pathogenesis, and likely exacerbates the observed mucinous phenotype.

Cross-species analyses demonstrated that a glycolysis signature is especially enriched in metaplastic epithelial cells in IPMNs. Besides the increased glycolysis (“Warburg effect”) generally observed in neoplastic cells [26], recent studies have suggested an intrinsic link between a glycolytic metabolism switch and somatic cell transdifferentiation and metaplasia. Such a glycolytic switch has been reported to be necessary in several contexts, including transdifferentiation of fibroblasts into induced endothelial cells [27], phenotypic switching of vascular smooth muscle cells [28], and of related interest, acinar-to-ductal metaplasia of the pancreas [29]. Our results suggest that glycolysis may be an accompaniment of mutant *GNAS*-mediated gastric (pyloric type) differentiation of neoplastic pancreatic epithelial cells. It is not fully understood why a glycolytic switch is observed in cells undergoing transdifferentiation or metaplasia. One possible reason is that reprogrammed cells need to adjust to new energy demands required for cell maintenance [30]. Epigenetic modifications needed to regulate gene expression during cell reprogramming may also play a role. Several glycolytic metabolites (e.g. pyruvate and lactate) can affect histone acetylation or lactylation [31]. Therefore, glycolysis may induce chromatin remodeling through these metabolites, which in turn affects transcription factor binding.

Our CRISPR/*Cas*9 loss-of-function screen and subsequent validation experiments demonstrated that glycolysis is an actionable vulnerability in *KRAS* and *GNAS* co-mutated cells. Whereas the relationship between glycolysis and IPMN has been largely unexplored until now, our findings suggest that mutant GNAS creates a targetable glycolysis dependency in the pathogenesis of IPMNs. As previously shown, glucose transport and glycolytic flux are required for growth of PDAC cells with oncogenic Ras [19]. The present study demonstrated that this requirement is significantly enhanced upon induction of mutant *GNAS* on a mutant *Ras* background. Increased glycolysis is generally related with tumor progression and aggressiveness via increased substrate production [32]. Lactate, which is the final product of glycolysis, can acidify the tumor microenvironment [32, 33]. Tumor microenvironment acidification promotes proliferation, resistance to apoptosis, invasiveness, metastatic potential, and aggressiveness of tumor cells [34]. Acidification of the neoplastic microenvironment may also modulate antitumor immunity through the attenuation of the activity and proliferation of T cells, which we have previously described accompanies IPMN progression [35] highlighting another avenue through which mutant *GNAS*-driven glycolysis may impact IPMN progression.

We identified the cAMP-PKA-PFKFB3 axis as a mechanism of increased glycolysis in *GNAS*-mutant metaplastic cells. One of the critical rate-limiting steps of glycolysis is the conversion of fructose-6-phosphate (F6P) to fructose-1,6-bisphosphate (F1,6P2), which is mediated by 6-phosphofructo-1-kinase (PFK-1). PFKFB is an enzyme that regulates the intracellular steady-state concentration of fructose 2,6-bisphosphate (F2,6P2), which is an activator of PFK-1 [36, 37, 38]. Among 4 isozymes (PFKFB1-4), PFKFB3 has the highest kinase:phosphatase activity ratio and is the most potent isozyme to enhance glycolysis, whose activity is regulated both at the transcriptional and post-transcriptional levels [36, 37, 38]. Interestingly, PFKFB3 is upregulated by mutant *KRAS*, which is well described to enhance glycolysis [39]. We found that PFKFB3 was activated through PKA-mediated phosphorylation, downstream of mutant *GNAS*. While *GNAS* mutations are observed in several tumor types other than IPMN-derived PDAC, such as colorectal cancer, lung adenocarcinoma, thyroid carcinoma, or pituitary adenoma [40, 41], the relationship between *GNAS* mutations and glycolysis has not been well described. Our findings may provide an insight into novel metabolic characteristics across multiple *GNAS*-mutant tumor types beyond the pancreas.

In conclusion, multimodal transcriptional analyses and functional genomics followed by the validation experiments demonstrated that mutant *GNAS* reprograms *Kras*-mutant pancreatic epithelial cells toward gastric (pyloric type) metaplasia in the pathogenesis of IPMNs, where increased glycolysis is essential for their maintenance. This epithelial reprogramming could enable us to understand the biology of the most common cystic precursor of PDAC, for which no specific therapeutics are currently available.

## Supporting information

Supplemental Files Immunoblots

## Acknowledgements

The authors acknowledge Drs. Bidyut Ghosh and Paola A Guerrero. The authors also thank Shui Ping So for technical assistance.

