## Supplementary figures and images for "Metabolic Reprogramming by Mutant GNAS Creates an Actionable Dependency in Intraductal Papillary Mucinous Neoplasms of the Pancreas"

### Supplemental figures

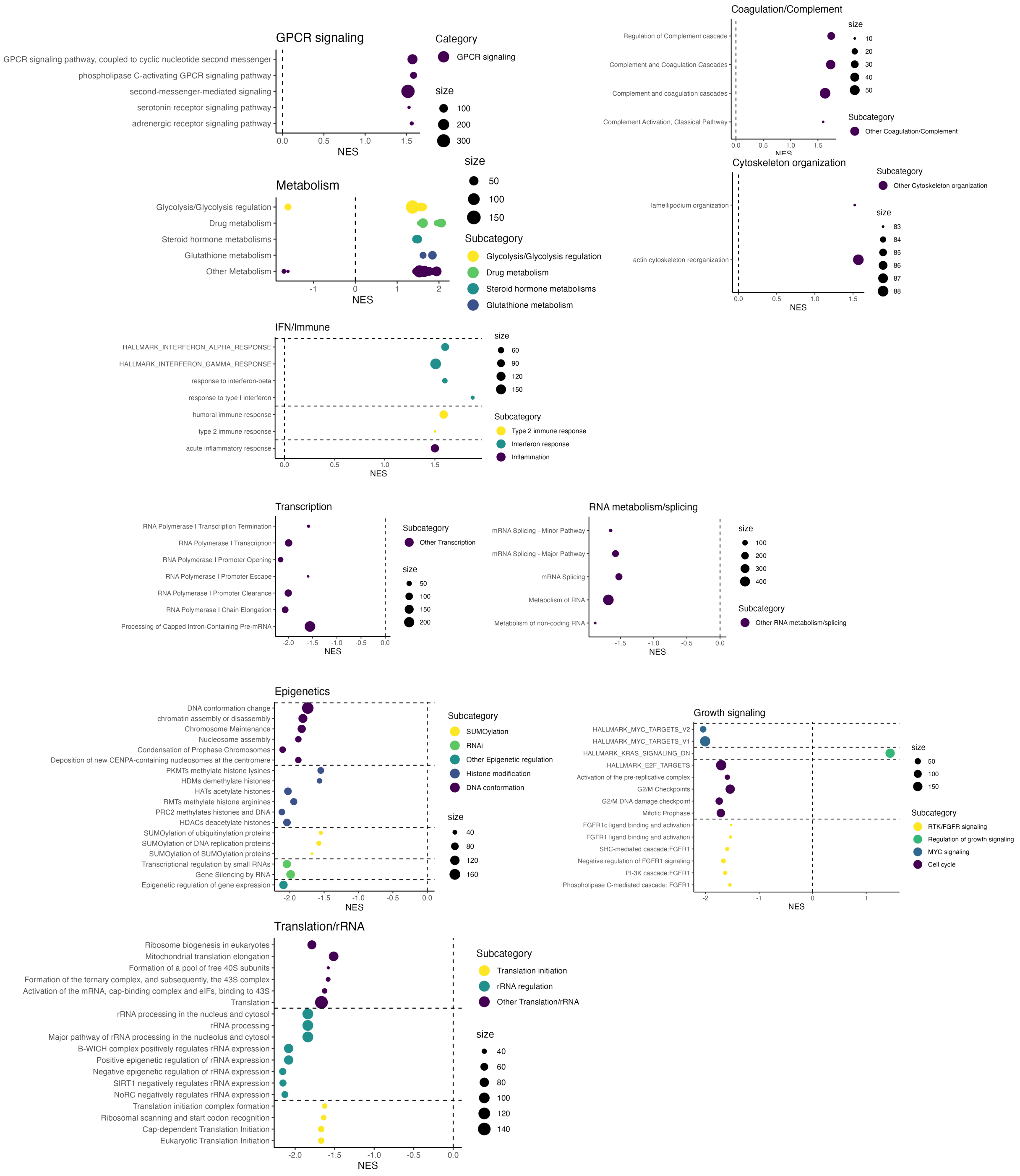


Supplementary Figure 1

(A)

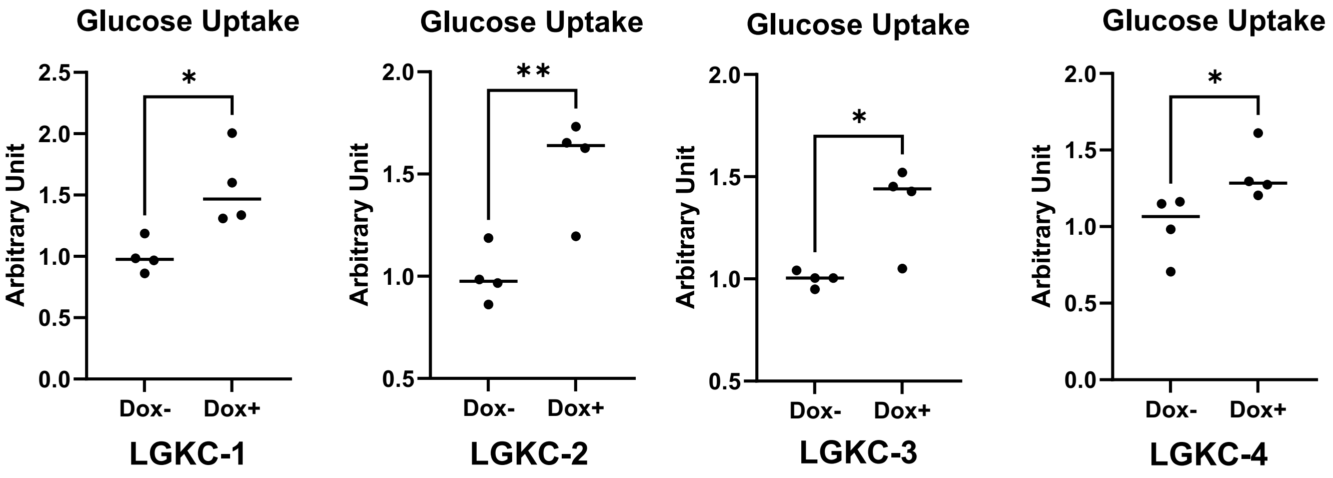


(B)

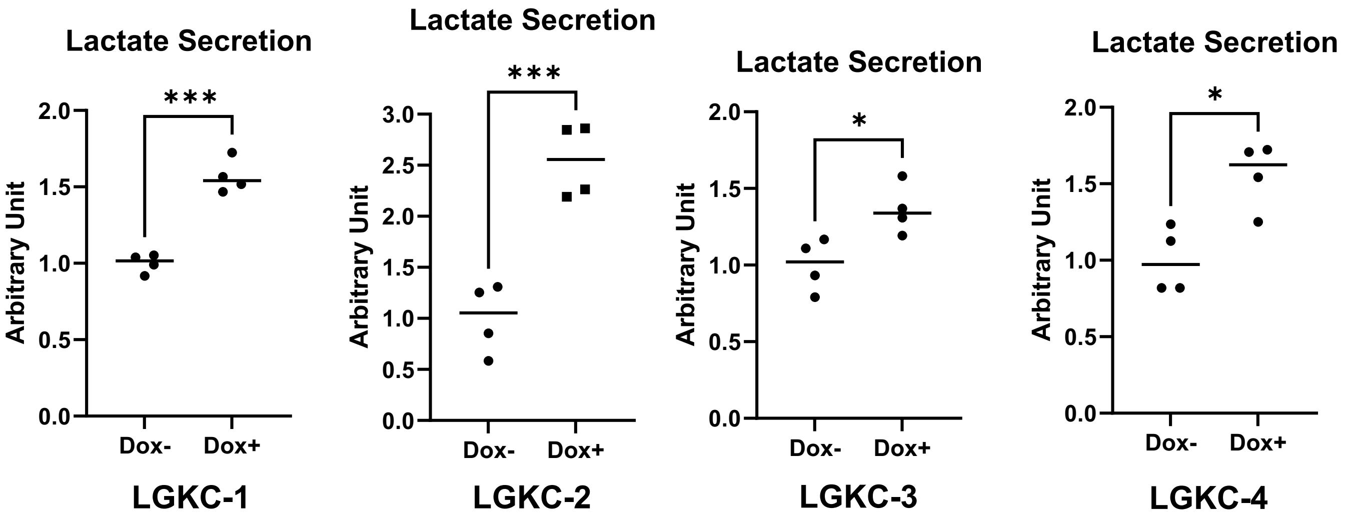


Supplementary Figure 2

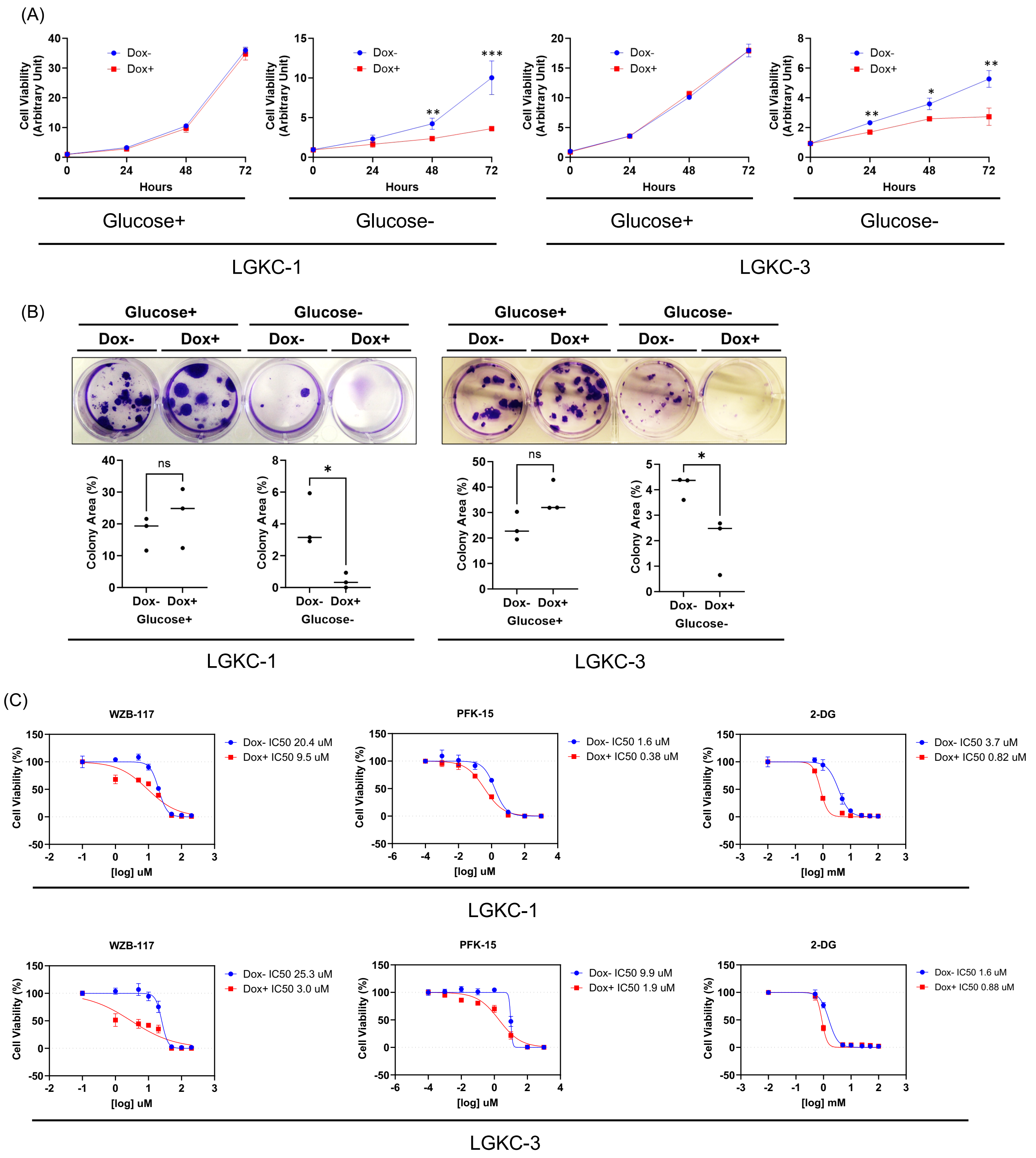


Supplementary Figure 3

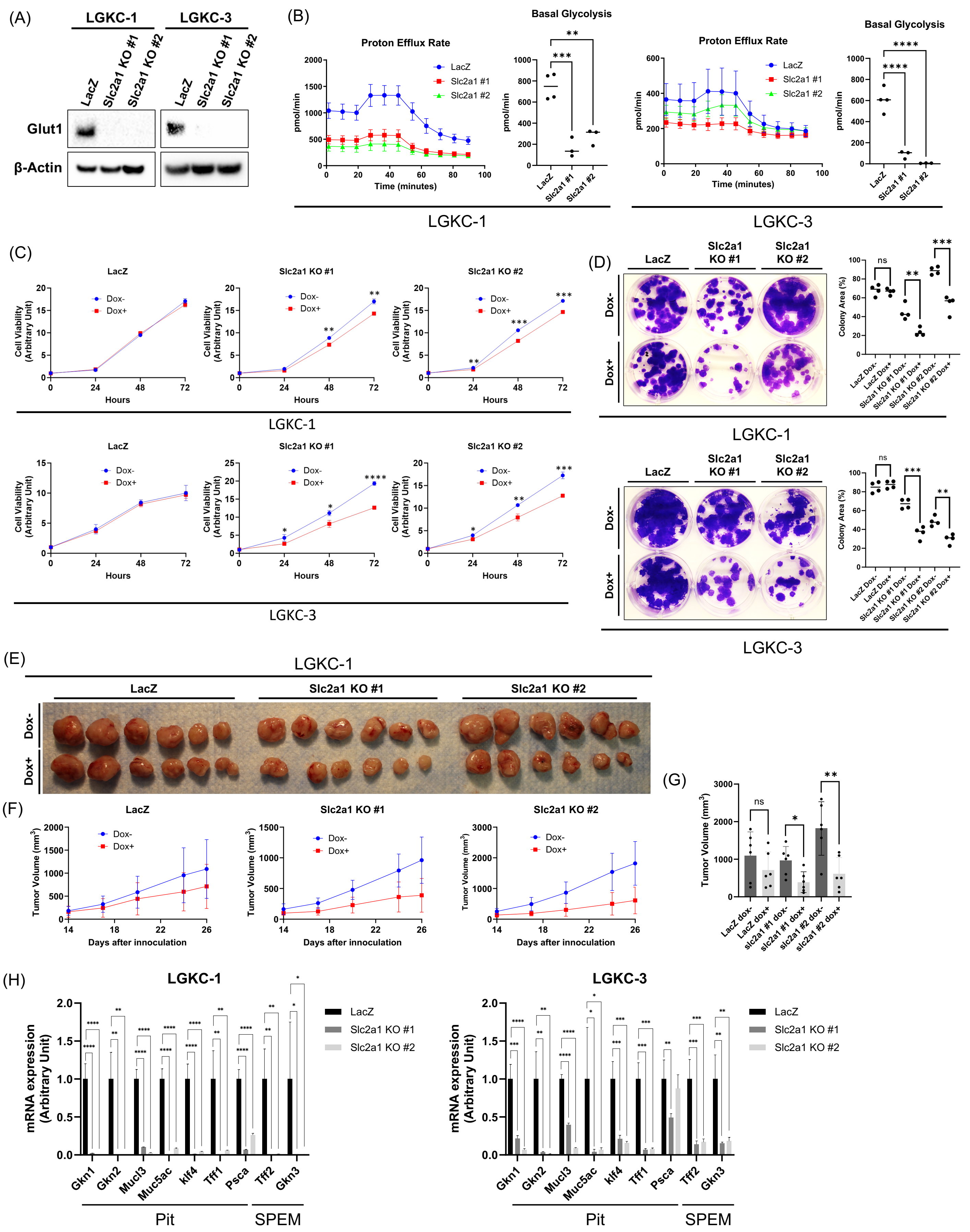


Supplementary Figure 4
