## Supplemental materials and methods for "Metabolic Reprogramming by Mutant GNAS Creates an Actionable Dependency in Intraductal Papillary Mucinous Neoplasms of the Pancreas"

**Table of contents**

Supplementary Materials and Methods: …………….……………2-9

Supplementary Figure legends: ……………….…………………..10-12

Supplementary Tables: ………………..……………………………13-15

References: ………………..………………………………………16-18

### Supplementary Materials and Methods

*Genome-wide CRISPR/Cas9 loss of function screening*

Mouse Toronto KnockOut version 3.0 (TKOv3) CRISPR lentiviral library was a kind gift from Moffat lab (University of Toronto) which contains 95,000 guides (5 guides/gene) targeting 19,000 genes. To generate lentivirus, HEK293T cells were transfected with mouseTKOv3, psPax2 (Addgene, Watertown, MA, USA), and pMD2.G plasmids (Addgene). For the transduction, LGKC-1 and LGKC-3 cells were incubated with the lentivirus for 48 hours. Following the puromycin selection for 48 hours, cells were re-seeded and divided into 6 groups (with or without 1 ug/mL doxycycline, in triplicate). Cells were trypsinized and merged every 3 days. After 9 days of incubation, genomic DNA was extracted using QIAamp DNA Blood Maxi Kit (QIAGEN, Hilden, Germany). Enrichment PCR was performed using KAPA HiFi HotStart ReadyMix PCR Kit (Roche, Basel, Switzerland) with the following primers (Forward; AGGGCCTATTTCCCATGATTCCTT, Reverse; TCAAAAAAGCACCGACTCGG). Indexing PCR was performed using the same reagent with unique pair of i5 and i7 indexing primers. Amplicons were extracted using E-gel and analyzed by Bioanalyzer. Next generation sequencing was performed by the Sequencing and Non-Coding RNA Program, MD Anderson Cancer Center. Raw reads from next generation sequencing were aligned to the TKOv3 mouse library using bowtie [2] to obtain read counts for all samples. The sgRNA reads were then processed with our BAGEL (Bayesian Analysis of Gene Essentiality) algorithm [3] to calculate the fold change at different timepoints compared to the initial time point of its corresponding screen and to calculate gene-level fitness scores (log BF). Positive BFs indicate higher confidence of gene essentiality, and negative scores indicate no fitness effect from knocking out that gene. DrugZ analysis [4] was also performed to compare doxycycline treated and untreated cells in each cell line to generate gene-level normZ scores with statistical significance. The genes were then ranked by their normZ scores to reveal synergistic and suppressor interactions.

*Cell culture experiments and reagents*

Glucose-free RPMI (Thermo Fisher Scientific, Waltham, MA, USA) supplemented with 10% dialyzed FBS (Thermo Fisher Scientific) and 1% penicillin and streptomycin was used for glucose deprivation. Cell viability was measured by WST-8 assay (Nacalai Tesque, Kyoto, Japan). We measured lactate secretion from cells into culture media using Lactate-Glo™ Assay (Promega, Madison, WI, USA), according to the manufacturer's protocol. Glucose Uptake-Glo™ Assay (Promega) was used to measure glucose uptake into cells. Forskolin and H-89 were purchased from Sigma-Aldrich (St. Louis, MO, USA) and Cayman Chemical (Ann Arbor, MI, USA), respectively. PFK-15, WZB-117, and 2-DG were purchased from MedChemExpress (Monmouth Junction, NJ, USA).

*Generation of CRISPR/Cas9-mediated knockout cells*

Guide RNAs (gRNAs) for *Gpi1*, *Slc2a1*, or control *LacZ* were cloned into lentiCRISPR v2 plasmid (Addgene). Detailed information of gRNA sequences is described in **Supplementary Table 1**. To generate lentivirus, Lenti-X 293T cells were transfected with the lentiCRISPR v2 with *Gpi1*, *Slc2a1*, or *LacZ* gRNA, psPax2 (Addgene), and pMD2.G plasmids (Addgene). LGKC cells were infected with the lentivirus, followed by puromycin (ThermoFisher Scientific) selection. Single clones were established by limiting dilution.

*Western blotting*

Cell pellet was lysed in M-PER Lysis Buffer (Thermo Fisher Scientific) supplemented with protease and phosphatase inhibitor (Sigma-Aldrich). Protein samples were prepared using NuPAGELDS Sample Buffer (Thermo Fisher Scientific) and NuPAGE Sample Reducing Agent (Thermo Fisher Scientific). Protein samples were separated by electrophoresis using NuPAGE® Bis-Tris gels (Thermo Fisher Scientific) and transferred onto a nitrocellulose membrane. For immunodetection, following antibodies were used: anti-Gpi1 (Thermo Fisher Scientific), anti-phospho-PFKFB3 (Thermo Fisher Scientific), anti-PFKFB3 (Proteintech, Rosemont, IL, USA), anti-β-actin (Santa Cruz Biotechnology, Dallas, TX, USA), anti-Glut1 (Abcam, Cambridge, UK), anti-phospho-PKA substrate (Cell Signaling Technology, Danvers, MA, USA), anti-HK1 (Cell Signaling Technology), anti-HK2 (Cell Signaling Technology), anti-LDHA (Cell Signaling Technology), anti-GAPDH (Cell Signaling Technology), anti-PFKP (Cell Signaling Technology), anti-PKM2 (Cell Signaling Technology), anti-pyruvate dehydrogenase (Cell Signaling Technology), anti-phospho-PFKFB2 (Cell Signaling Technology), anti-PFKFB2 (Cell Signaling Technology), and anti-PFKFB4 (Thermo Fisher Scientific). **The entire compendium of whole gel blot images is included in the Supplementary Data file**.

*Quantitative real-time PCR*

RNeasy Mini Kit (QIAGEN, Hilden, Germany) was used to extract total RNA according to the manufacturer's protocol. cDNA was subsequently produced by reverse transcription using iScript™ Reverse Transcription Supermix for RT-qPCR (Bio-Rad, Hercules, CA). Real-time RT-PCR was performed to measure gene expression using the primers shown in **Supplementary table 2** purchased from Integrated DNA Technologies (San Jose, CA, USA). All gene expression levels were normalized to those of β-actin and relative expression was calculated using the ΔΔCt method.

*Ex vivo metabolic analysis*

Seahorse XFe24 Extracellular Flux Analyzer (Agilent, Santa Clara, CA, USA) was used to measure Oxygen Consumption Rates (OCR) and Extracellular Acidification Rate (EACR) of cells in real-time [5]. We used Seahorse XF Glycolytic Rate Assay kit (Agilent, Santa Clara, CA, USA) according to the manufacturer's instructions.

*Live cell studies with $^{13}C$ hyperpolarized pyruvate MRS*

These studies were conducted as we have previously described [6]. Briefly, Ox063 trityl radical (GE Healthcare) was mixed with neat [1-$^{13}$C] pyruvic acid (Sigma-Aldrich, St. Louis, MO, USA), to a concentration of 15 mmol/L. 25 uL of this solution with 0.4uL of 1.5mM chelated gadolinium (Magnevist, Bayer Healthcare Wayne, NJ, USA) was loaded into a DNP commercial HyperSense polarizer (Oxford Instruments) and irradiated at a microwave frequency of 94.100 GHz for 45 minutes (until polarization plateau) and then dissolved in 4 mL buffer solution containing 40 mmol/L Tris (7.6 pH preset), 80 mmol/L NaOH, 0.1 g/L EDTA, and 50 mmol/L NaCl. All cell lines experiments were performed on a vertical bore 7T Bruker BioSpin NMR with 10 mm broadband probe and running TopSpin3.5 software. Twenty million cells were suspended in 2.0 mL RPMI cell culture media (with 10% D2O) and kept in a 10-mm NMR tube. The NMR probe was heated to 37°C and was maintained at this temperature throughout the experiment. After auto-locking, tuning, and shimming were performed, 500 uL of the hyperpolarized [1-$^{13}$C] pyruvate was injected into the suspended cells using an injection port. The final concentration of hyperpolarized pyruvate in the NMR reactor was approximately 8.7 mmol/L. Single $^{13}$C transients were taken using Waltz decoupling (zgdc pulse) every 6 seconds with a 15° flip angle. The lactate / pyruvate ratio was calculated as the total lactate signal divided by the total pyruvate signal over the first 3 minutes of the experiment. Cell viability was analyzed using trypan blue staining and counted on a BioRad TC20 automated cell counter. The percentages of viable cells were similar before and after the experiments (90%–95% viable). The lactate / pyruvate ratio was normalized as per million cells times 1000.

*In vivo animal studies with $^{13}$C hyperpolarized pyruvate MRS*

We performed $^{13}$C pyruvate Hyperpolarozed MRI to evaluate glycolysis flux i*n vivo* as previously reported [6, 7]. From 8 weeks of age, *Kras;Gnas* mice were randomly assigned to either doxycycline or normal diets (control group) for up to 15 weeks. Six mice were used per group and all mice were included in the analysis. $^{13}$C Pyruvate was hyperpolarized as above. After dissolution, 80 mmol/L hyperpolarized 1-$^{13}$C pyruvate solution with physiological pH 7.6 was injected into each mouse via a tail vein catheter. All imaging and spectroscopy were performed with a dual tuned $^{1}H/^{13}C$ volume coil (OD: 75 mm, ID: 40 mm, Bruker BioSpin MRI GmbH) in a 7T Bruker Biospec horizontal bore MR scanner (Bruker BioSpin MRI GmbH) equipped with a single channel for carbon excitation/reception. Proton anatomic images were taken using a multislice T2-weighted RARE sequence. A small 8 mol/L $^{13}C$-urea phantom doped with gadolinium-DPTA was placed on abdominal site of the mouse (head-supine position) to allow for chemical shift referencing. For $^{13}C$ spectroscopy, a series of slice-selective $^{13}C$ spectra (field-of-view 40x40 $mm^2$, slice-thickness 10 mm) were collected right after injection of hyperpolarized pyruvate. A total of 90 transients were acquired at 2-second intervals (total time 3 minutes). Each transient utilized a 20° flip angle excitation pulse (gauss pulse) and 2,048 data points. Data were processed by MATLAB (MathWorks Inc.), TopSpin (Bruker BioSpin GmbH), and MestReNova (Mestrelab Research) software. The dynamic spectra were manually phased, and line broadening was applied (10 to 15 Hz). The areas under the spectral peaks for pyruvate and lactate were integrated over the whole array. Lactate / pyruvate ratio was calculated as total lactate signal divided by total pyruvate signal. The sample size was determined using G*Power (3.1.9.7) assuming an effect size of 2, alpha = 0.05 and 80% power (beta = 0.2). n refers to individual mice and statistical analysis was performed with Student's t-test in Prism Prism 10 (GraphPad Software, Inc.).

*Colony formation assay*

The protocol published previously was modified for optimization [8]. Briefly, 500 cells per well were plated in 12-well plates. Colonies were fixed and stained 10 days after incubation with or without doxycycline. Colony area per well was quantified using ImageJ software (NIH, Bethesda, MD, USA).

*Immunohistochemistry*

Immunohistochemical staining was performed using paraffin-embedded sections of the pancreas obtained from human IPMN samples (7 *GNAS*-wild type and 7 *GNAS*-mutant), which have been used for our previous report as part of the NCI Cancer Moonshot Precancer Atlas Pilot Project (PCAPP) [9]. An anti-phospho-PFKFB3 antibody was purchased from Thermo Fisher Scientific. Color development was performed by using diaminobenzidine tetrahydrochloride (DAB) solution. Staining score was classified into 4 categories as "Negative", "Weak", "Moderate", and "Strong".

*Subcutaneous injection model*

Eight-week-old female athymic mice (NU/J, #002019) for transplantation experiments were purchased from The Jackson Laboratory (Bar Harbor, ME, USA). Mice were subcutaneously inoculated in the bilateral flank portions with $1 \times 10^6$ *Gpi1* KO (clone #1 or clone #2), *Slc2a1* KO (clone #1 or clone #2) or *LacZ* (2 control groups) cells (6 mice per group) suspended in Matrigel (Corning). The length and width of the subcutaneous tumors were measured every 3 to 4 days. Tumor volume was calculated according to the following formula: tumor volume = $1/2 \times (\text{major axis}) \times (\text{minor axis})^2$ [10, 11]. The transplanted mice were randomly assigned to doxycycline or standard feed (control group) using the coin toss method. The experiment was not blinded. A total of 18 mice were used.9 The number of mice was selected based on prior experience. n refers to individual tumors (2 bilateral tumors per mouse). All transplanted animals were included in the analysis. Statistical differences were determined with Student's t-test using Prism 10 (GraphPad Software, Inc.).

*RNA-sequencing and data analysis*

*Kras;Gnas* cells were treated with 1 ug/mL doxycycline or vehicle for 24 hours before RNA collection. Total RNA was extracted from LGKC cells using RNeasy Mini Kit (QIAGEN) according to the manufacturer's protocol. 76 bp paired-end stranded RNA sequencing was performed on a HiSeq 4000 sequencer (Illumina, CA, USA). Library preparation and sequencing was performed by the MD Anderson Sequencing and Microarray Facility (SMF). Raw reads from sequencing were quantified with featureCounts [12] after mapping to Mus musculus genome (UCSC mm10) genome with STAR [13].

Genes significantly expressed in doxycycline-induced samples were identified using pairwise (doxycycline vs vehicle) analysis with DESeq2 [14]. GSEA [15] (http://software.broadinstitute.org/gsea/index.jsp) and Webgestalt [16] software packages were used to find enriched gene sets using the HALLMARK, REACTOME, KEGG and Gene Ontology – Biological Process (GO-BP) databases. Specific gene sets of interest were downloaded from the MSigDB database (https://www.gsea-msigdb.org/gsea/msigdb/index.jsp). Significantly enriched gene sets were manually curated into categories and subcategories based on similarities and shared search terms.

*Single cell RNA-sequencing (scRNA-seq) and data analysis*

The detailed protocol for murine pancreatic scRNA-seq has been described by us previously [17, 18]. Briefly, pancreas tissues were enzymatically digested into a single-cell suspension. scRNAseq library was generated by 10X Chromium System (10X Genomics Inc, Pleasanton, CA). Samples were run on the Illumina NextSeq500 High Output Flowcell (Illumina). For data analysis, we used Cell Ranger, version 3.0.1 (10X Genomics), to process raw sequencing data. After sequencing and data processing, downstream analyses were performed using Seurat (v4.3.0) [19] package in R (v4.0.0). Cells with unique feature counts over 5000 or less than 200 and with >25% mitochondrial counts were filtered out as quality control measure. Individual samples were integrated with Harmony (v 0.1.1) [20] package within R to remove the batch effects. Single cells were clustered with FindNeighbors and FindClusters functions within Seurat and they were annotated to know cell types using SingleR [21] and known gene markers. Rank-based single cell GSEA was performed using the irGSEA 2.1.5 package.

*Spatial transcriptomics and data analysis*

The detailed protocol for human and murine pancreatic spatial transcriptomics has been described by us previously [22]. Briefly, 7 low-grade IPMN samples were used as patient samples for analysis. All biospecimens were collected under IRB approved protocols and obtained using written informed consent.

At MDACC, tissue samples were collected under the Pancreas Tissue Bank, LAB00-396 and subsequently analyzed under a use protocol PA15-0014. Pancreata collected from *Kras;Gnas* mice at 33 weeks of age (25 weeks of doxycycline diet or normal diet) were used for murine samples. Samples were cryosectioned onto Visium Spatial Gene Expression slides (10X Genomics), followed by library preparation and sequencing using a NextSeq 500 (Illumina). Sequencing data were preprocessed with SpaceRanger (v.1.2.1) while mapping to the latest human (hg38) or mouse (mm10) reference genome. Filtered matrices were imported into R with Seurat package for downstream analysis. As stated previously [22], histological annotation of areas within human and mouse spatial transcriptomics data were manually conducted using the Loupe Browser. Robust cell type deconvolution (RCTD) algorithm [23] was used to assess the cell type proportions in each spot.

*Statistical analysis*

Student's t-test was used to compare differences between unpaired or paired groups with normal distributions using Prism 10 (GraphPad Software, Inc.). Chi-square test was performed to compare categorical variables. Pearson Correlation Coefficients were calculated using Prism 10 (GraphPad Software, Inc.) or in R. For comparisons with more than two groups, we used one-way ANOVA with multiple comparison correction in Prism 10 (GraphPad Software, Inc.). p-values less than 0.05 were considered statistically significant.

**Supplementary Figure legends**

**Supplementary Figure 1. Gene set enrichment analysis (GSEA) of RNA sequencing revealed drastic transcriptional change across multiple biological categories by the induction of mutant *GNAS* in *Kras;Gnas* cells.**

GSEA of subcategories from each biological category that showed positive or negative enrichment (presented in Figure 1C) is shown. Total RNA for RNA sequencing was collected after 24-hour incubation with or without 1 μg/mL doxycycline. Paired analysis was performed comparing doxycycline-positive over doxycycline-negative samples in 4 cell lines (LGKC-1, 2, 3, and 4).

**Supplementary Figure 2. Glucose uptake and lactate secretion assays in individual *Kras;Gnas* cell lines.**

Results of glucose uptake assay and lactate secretion assay in each *Kras;Gnas* cell line (presented in Figure 5A and 5B) are shown. LGKC-1, 2, 3, or 4 cells were incubated with or without 100 ng/mL doxycycline for 24 hours before each assay (N = 4, technical replicates).

(A) Glucose uptake assay.

(B) Lactate secretion assay.

*p<0.05, **p<0.01, ***p<0.001.

**Supplementary Figure 3. Glucose deprivation and inhibition suppressed proliferation of *Kras;Gnas* cells with *Gnas*$^{R201C}$ expression.**

(A-B) *Kras;Gnas* cells were plated with glucose-containing medium without doxycycline. Twenty-four hours after plating, the medium was replaced either with glucose-containing medium or glucose-free medium, with or without 100 ng/mL doxycycline.

(A) Cell proliferation assay. Cell viability was measured by WST-8 assay at indicated time points. (N = 3, technical replicates)

(B) Colony formation assay. Cells were stained after 10-day incubation with or without doxycycline.

(C) Dose-response curve and the half maximal inhibitory concentration ($IC_{50}$) for cell viability of *Kras;Gnas* cells under the treatment with glycolysis inhibitors. Cells were treated with WZB-117, PFK-15, or 2-Deoxy-D-glucose (2-DG) at indicated concentrations with or without 100 ng/mL doxycycline for 72 hours. N = 3, technical replicates.

*p<0.05, **p<0.01, ***p<0.001.

**Supplementary Figure 4. Loss of *Slc2a1* decreased glycolysis and suppressed the growth of *Kras;Gnas* cells with *Gnas*$^{R201C}$ expression both *in vitro* and *in vivo*.**

*Slc2a1* or control *lacZ* were knocked out by CRISPR/*Cas9* in *Kras;Gnas* cells.

(A) Western blotting of *Slc2a1* KO and control *Kras;Gnas* cells. Cells were treated with 100 ng/mL doxycycline for 24 hours before protein collection.

(B) Seahorse XF Glycolytic Rate Assay. Cells were incubated with 100 ng/mL doxycycline for 24 hours before the assay. In the assay, cells were incubated in the medium containing glucose, glutamine, and pyruvate, followed by the injection of Rotenone and Antimycin (Rot/AA) and 2-Deoxy-D-glucose (2-DG). Proton efflux rate (PER) was measured at each time point to calculate basal glycolysis.

(C) Cell proliferation assay. WST-8 assay was performed to measure cell viability at indicated time points. N = 3, technical replicates.

(D) Colony formation assay. Cells were stained after 10-day incubation with or without 100 ng/mL doxycycline.

(E-G) Allograft injection model of *Slc2a1* KO cells. Control (*LacZ*) or *Slc2a1* KO *Kras;Gnas* cells were subcutaneously inoculated to the bilateral flank portion of nude mice (N = 6 per group). Mice were fed with normal diet or doxycycline diet from the day of inoculation. Tumor volumes were measured every 2 to 4 days from day 14 until sacrifice at day 26.

(E) Macroscopic appearance of the tumors.

(F) Sequential tumor volume as a function of time after cell inoculation.

(G) Tumor volume at the time of sacrifice (day 26).

(H) Quantitative PCR analysis for gastric pit cell and spasmolytic polypeptide expressing metaplasia (SPEM) markers in *Slc2a1*-KO *Kras;Gnas* cells. Cells were incubated with 100 ng/mL doxycycline for 24 hours before RNA collection. N = 4, technical replicates.

*p<0.05, **p<0.01, ***p<0.001, ****p<0.0001.

**Supplementary Table 1. Sequence of gRNAs.**

| Name | Sequence | Supplier |
|---|---|---|
| gpi1 gRNA #1 | CAACTTCAGGTGCACGCACG | IDT |
| gpi1 gRNA #2 | AACAACTTCAGGTGCACGCA | IDT |
| slc2a1 gRNA #1 | CGGCTCCGGTATCGTCAACA | IDT |
| slc2a1 gRNA #2 | GTTGACGATACCGGAGCCGA | IDT |
| lacZ gRNA | CCCGAATCTCTATCGTGCGG | IDT |

**Supplementary Table 2. Sequence of primers for quantitative PCR.**

| Name | Sequence | Supplier |
|---|---|---|
| Mouse gkn1 Fw | CCGCCATGAAGCTCACAATG | IDT |
| Mouse gkn1 Rv | CCAGGCCCTTTACCCTTCTG | IDT |
| Mouse gkn2 Fw | AACATCCACTCAGGCTCGTG | IDT |
| Mouse gkn2 Rv | CTGTCGAATAGGCGACCCAA | IDT |
| Mouse gkn3 Fw | CTCCAGCTTCTGTCCTTCATGG | IDT |
| Mouse gkn3 Rv | AGACACTTCTCACAGGCAGAGG | IDT |
| Mouse mucl3 Fw | CCTGCTGGTTTTCATTGGCCTG | IDT |
| Mouse mucl3 Rv | CTGCTCCATCAGGTAAACTGGG | IDT |
| Mouse muc5ac Fw | TCTACTGACTGCACCAACACA | IDT |
| Mouse muc5ac Rv | CCACACTTTCGCAGCTCAAC | IDT |
| Mouse klf4 Fw | GAAATTCGCCCGCTCCGATGA | IDT |
| Mouse klf4 Rv | CTGTGTGTTTGCGGTAGTGCC | IDT |
| Mouse tff1 Fw | CCCGGGAGAGGATAAATTGT | IDT |
| Mouse tff1 Rv | GCCAGTTCTCTCAGGATGGA | IDT |
| Mouse tff2 Fw | TGCTTTGATCTTGGATGCTG | IDT |
| Mouse tff2 Rv | GGAAAAGCAGCAGTTTCGAC | IDT |
| Mouse psca Fw | GCACAGTTGCTTTACATCGCGC | IDT |

| | | |
|---|---|---|
| Mouse psca Rv | ACAGGTCAGAGTAGCAGCACGT | IDT |
| Mouse β-actin Fw | CACAGCTTCTTTGCAGCTCCTT | IDT |
| Mouseβ-actin Rv | CGTCATCCATGGCGAACTG | IDT |
