## Supplemental Files Immunoblots for "Metabolic Reprogramming by Mutant GNAS Creates an Actionable Dependency in Intraductal Papillary Mucinous Neoplasms of the Pancreas"

Figure 6A

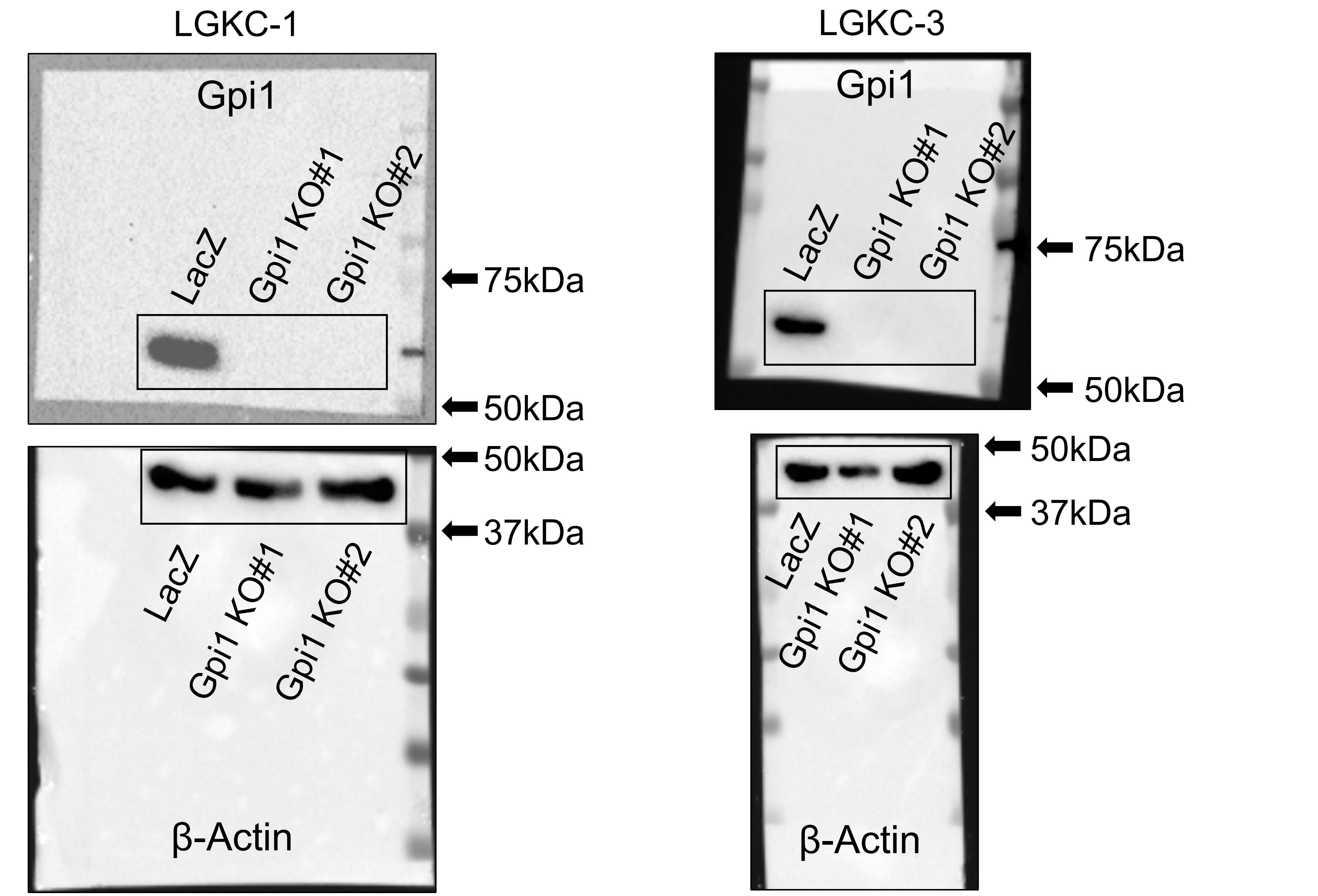
LGKC-1
Gpi1
LacZ
Gpi1 KO#1
Gpi1 KO#2
75kDa
50kDa
50kDa
37kDa
LacZ
Gpi1 KO#1
Gpi1 KO#2
β-Actin
LGKC-3
Gpi1
LacZ
Gpi1 KO#1
Gpi1 KO#2
75kDa
50kDa
50kDa
37kDa
LacZ
Gpi1 KO#1
Gpi1 KO#2
β-Actin

Figure 7A

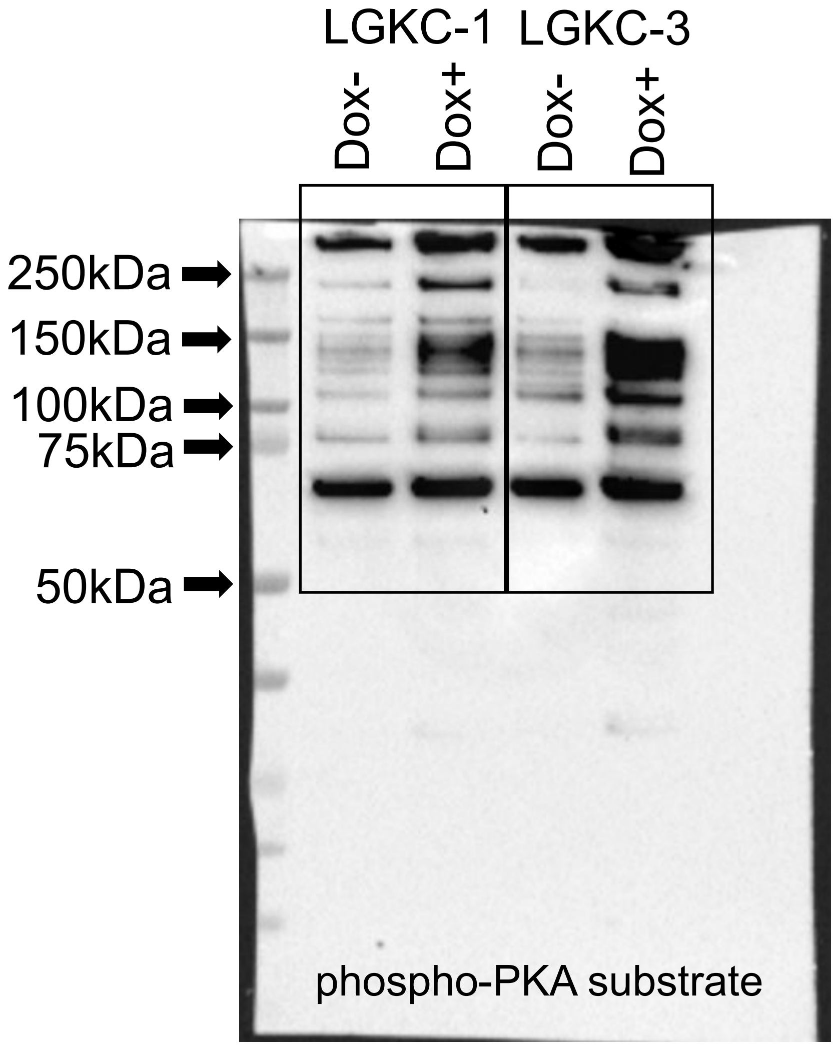
LGKC-1
LGKC-3
Dox-
Dox+
Dox-
Dox+
250kDa
150kDa
100kDa
75kDa
50kDa
phospho-PKA substrate

Figure 7A

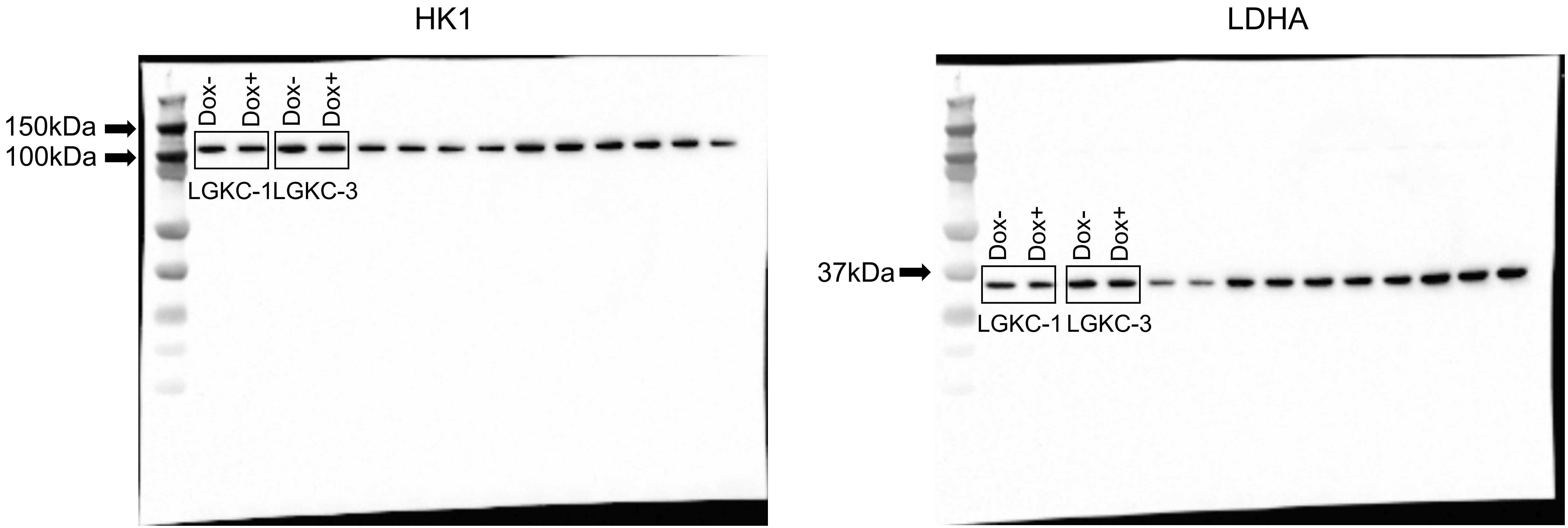
HK1
150kDa
100kDa
Dox-
Dox+
Dox-
Dox+
LGKC-1
LGKC-3
LDHA
37kDa
Dox-
Dox+
Dox-
Dox+
LGKC-1
LGKC-3

Figure 7A

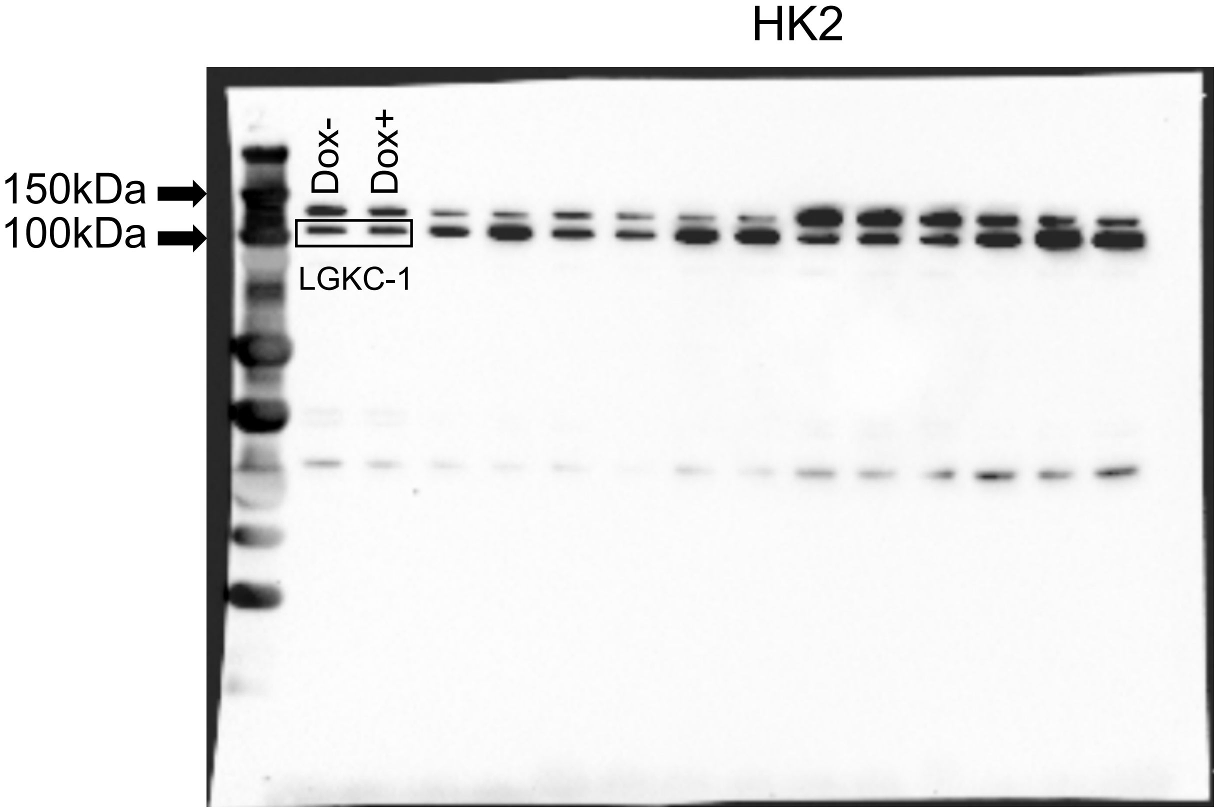
HK2
150kDa
100kDa
Dox-
Dox+
LGKC-1

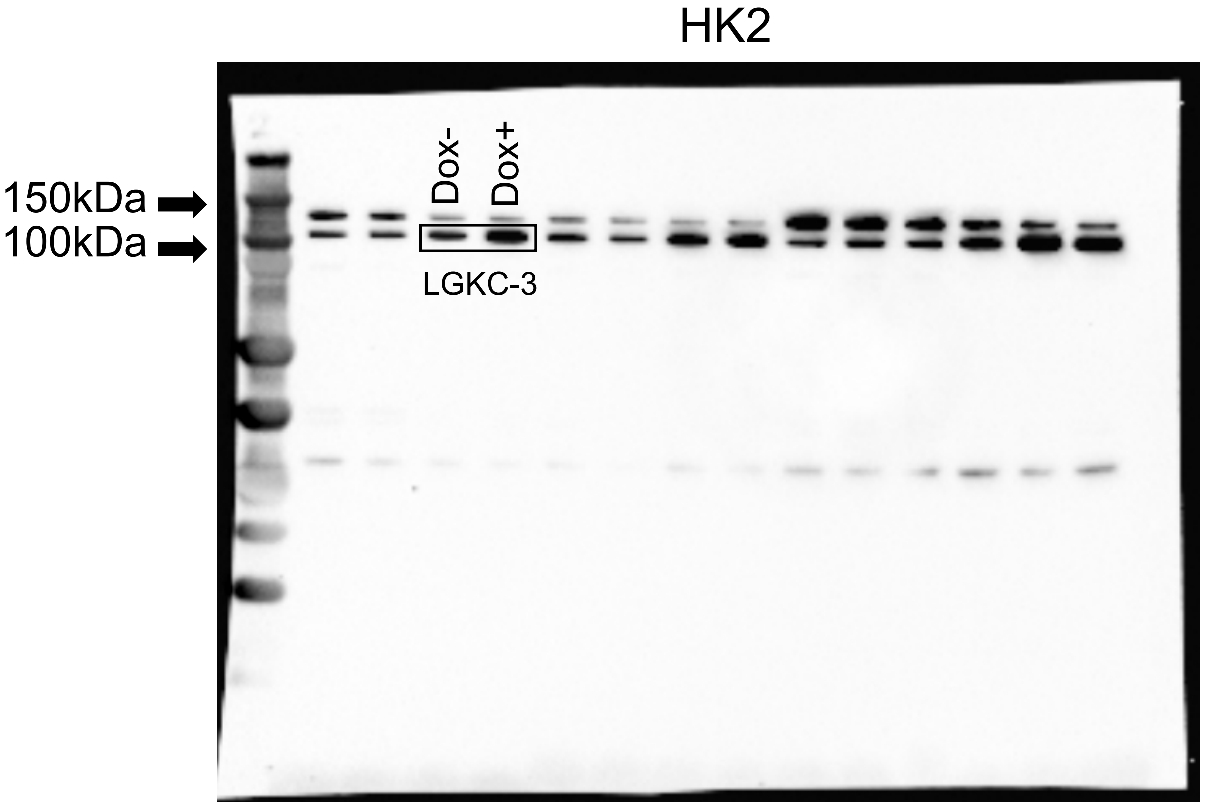
HK2
150kDa
100kDa
Dox-
Dox+
LGKC-3

Figure 7A

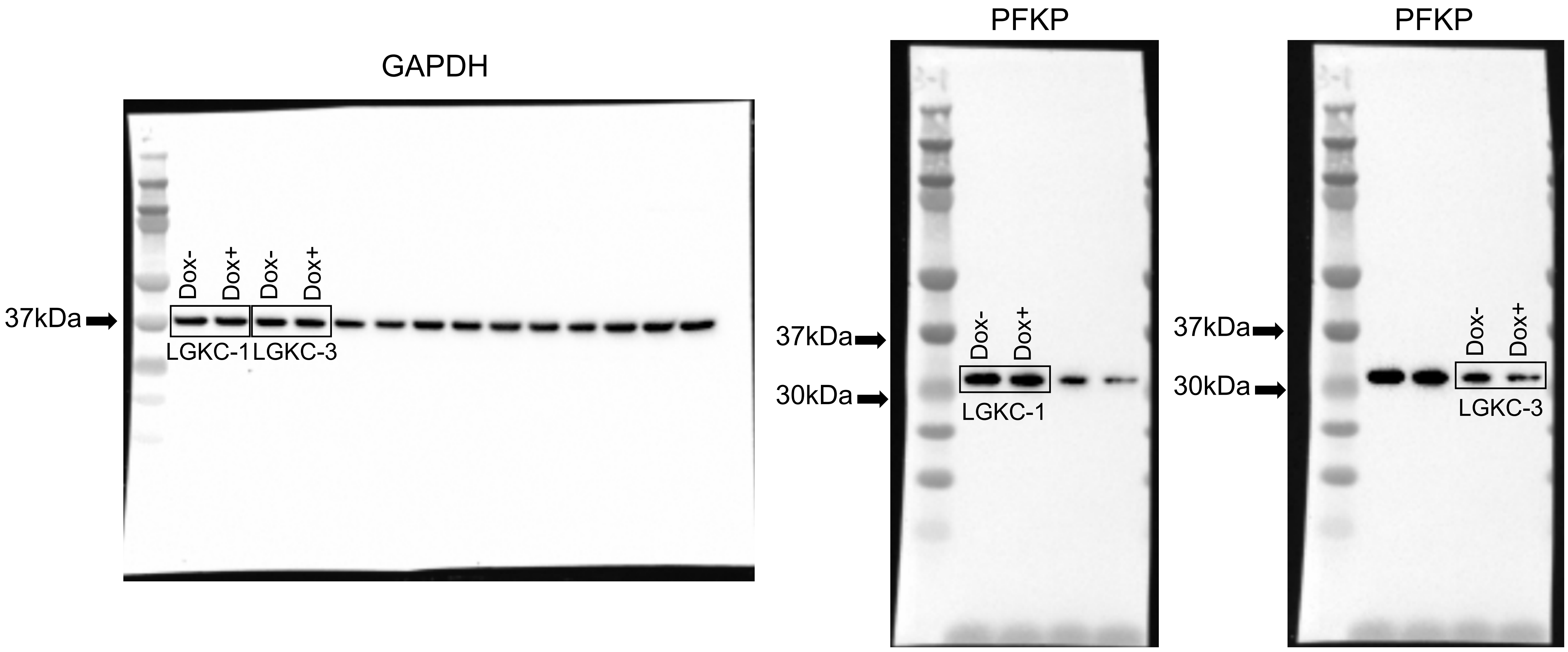

GAPDH
37kDa
Dox-
Dox+
Dox-
Dox+
LGKC-1
LGKC-3
PFKP
37kDa
30kDa
Dox-
Dox+
LGKC-1
PFKP
37kDa
30kDa
Dox-
Dox+
LGKC-3

Figure 7A

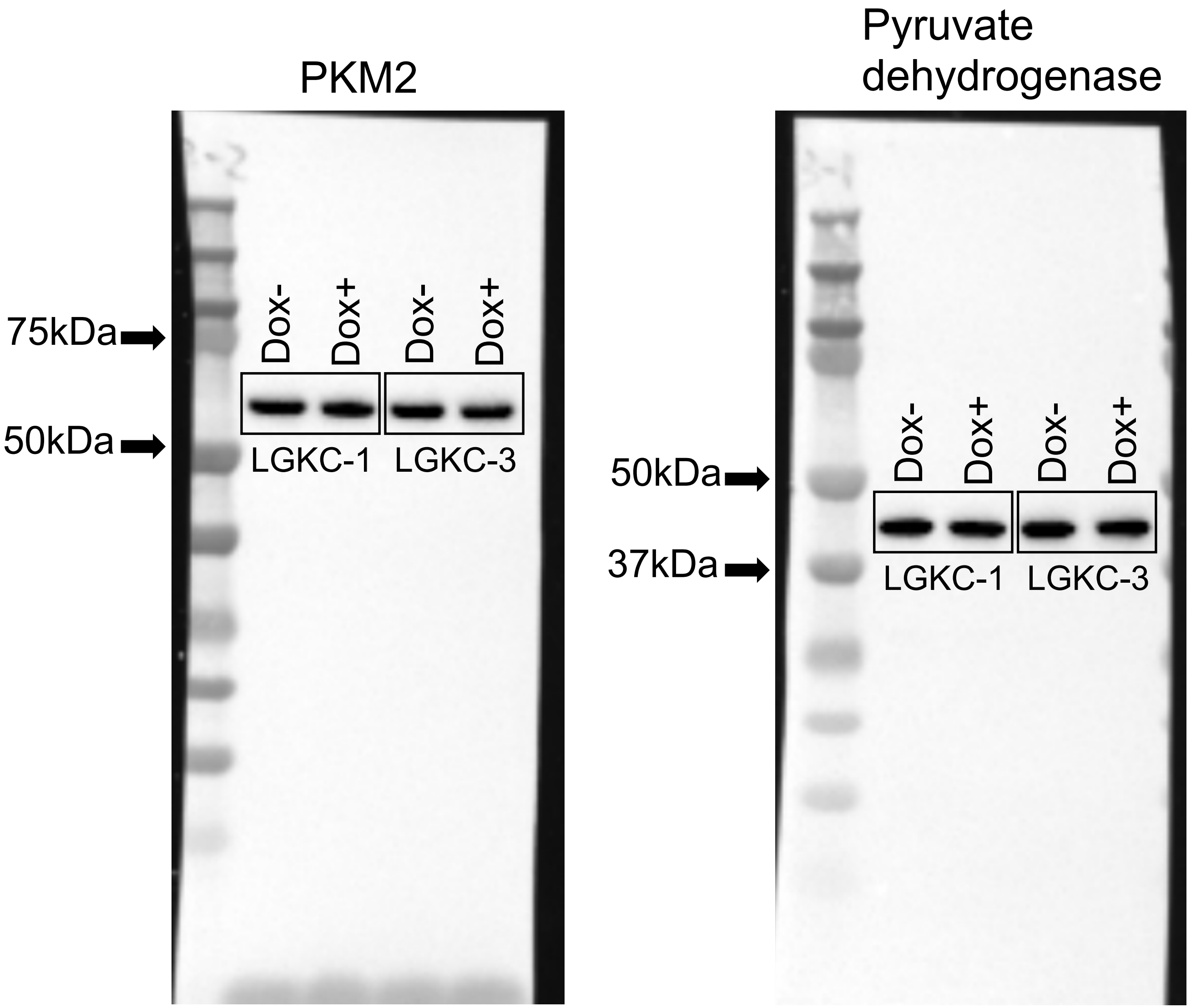
PKM2
75kDa
50kDa
Dox-
Dox+
Dox-
Dox+
LGKC-1
LGKC-3
Pyruvate dehydrogenase
50kDa
37kDa
Dox-
Dox+
Dox-
Dox+
LGKC-1
LGKC-3

Figure 7A

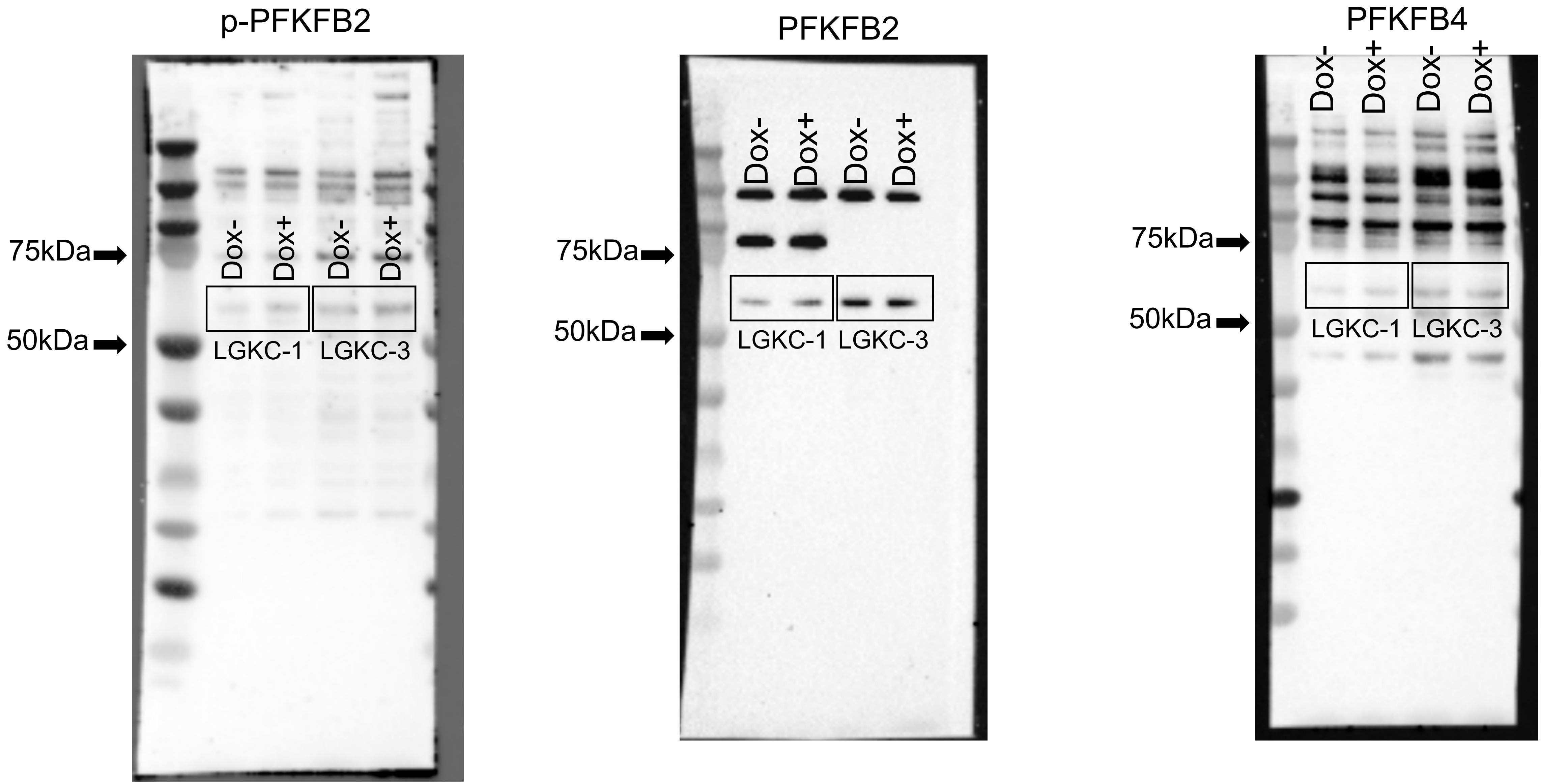
p-PFKFB2
PFKFB2
PFKFB4
Dox-
Dox+
Dox-
Dox+
75kDa
50kDa
LGKC-1
LGKC-3

Figure 7A

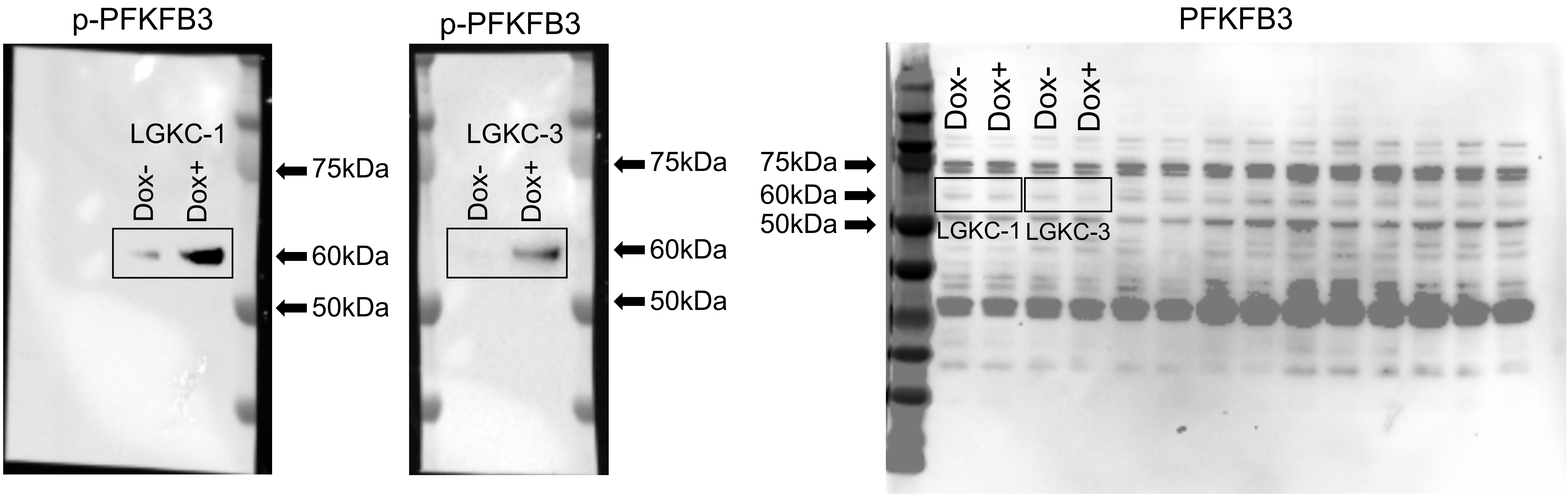
p-PFKFB3
LGKC-1
Dox-
Dox+
75kDa
60kDa
50kDa
p-PFKFB3
LGKC-3
Dox-
Dox+
75kDa
60kDa
50kDa
PFKFB3
Dox-
Dox+
Dox-
Dox+
75kDa
60kDa
50kDa
LGKC-1
LGKC-3

Figure 7B

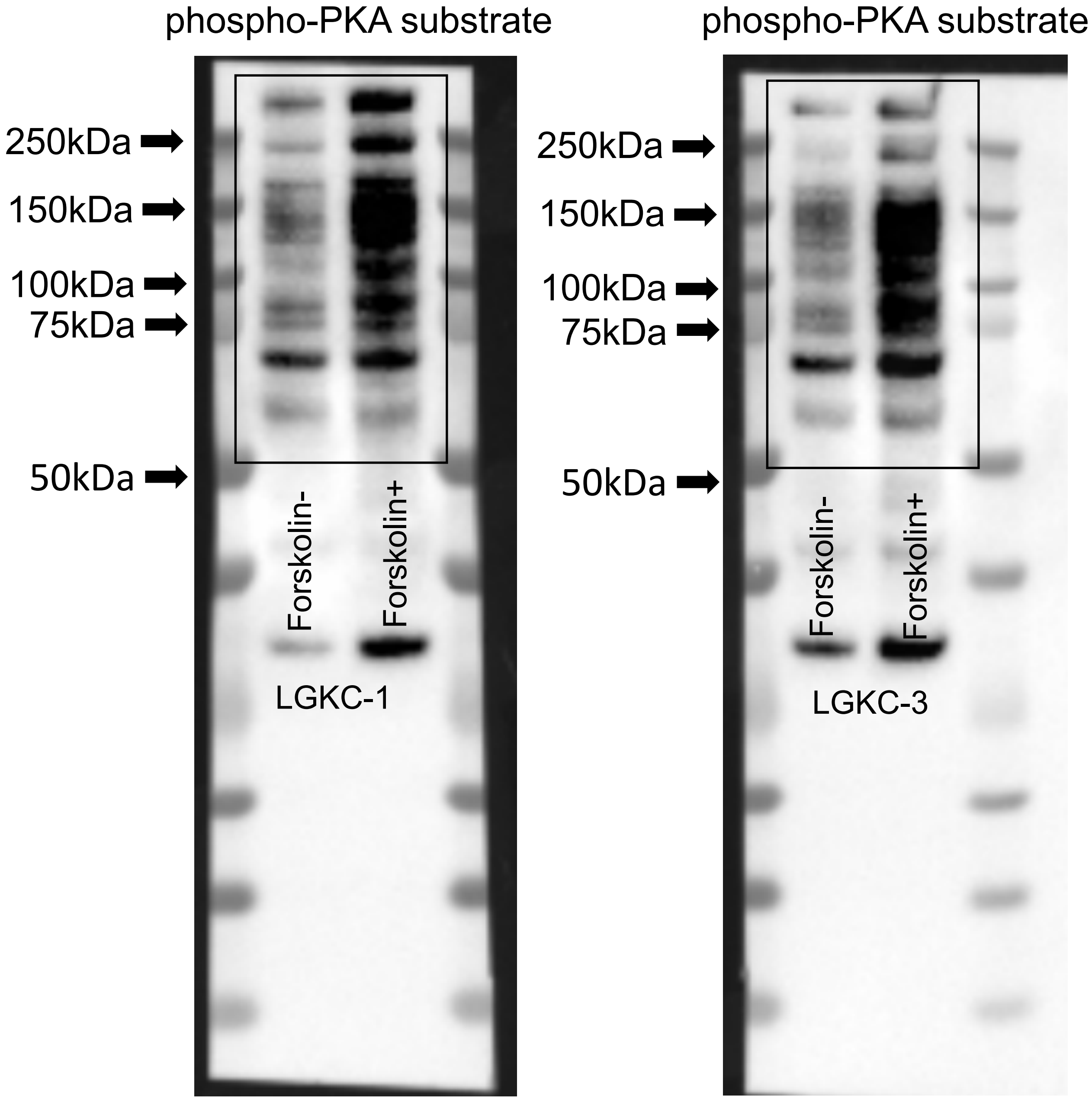
phospho-PKA substrate
250kDa
150kDa
100kDa
75kDa
50kDa
Forskolin-
Forskolin+
LGKC-1
phospho-PKA substrate
250kDa
150kDa
100kDa
75kDa
50kDa
Forskolin-
Forskolin+
LGKC-3

Figure 7B

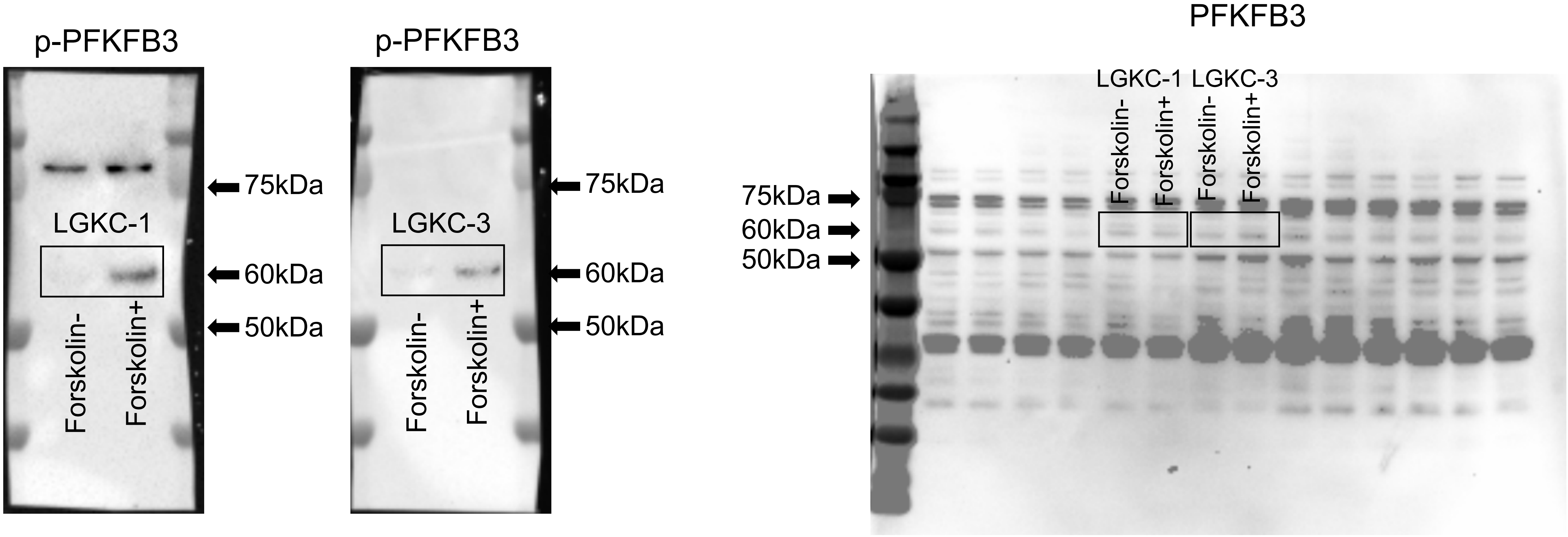
p-PFKFB3
75kDa
LGKC-1
60kDa
50kDa
Forskolin-
Forskolin+
p-PFKFB3
75kDa
LGKC-3
60kDa
50kDa
Forskolin-
Forskolin+
PFKFB3
LGKC-1
LGKC-3
Forskolin-
Forskolin+
Forskolin-
Forskolin+
75kDa
60kDa
50kDa

Figure 7B

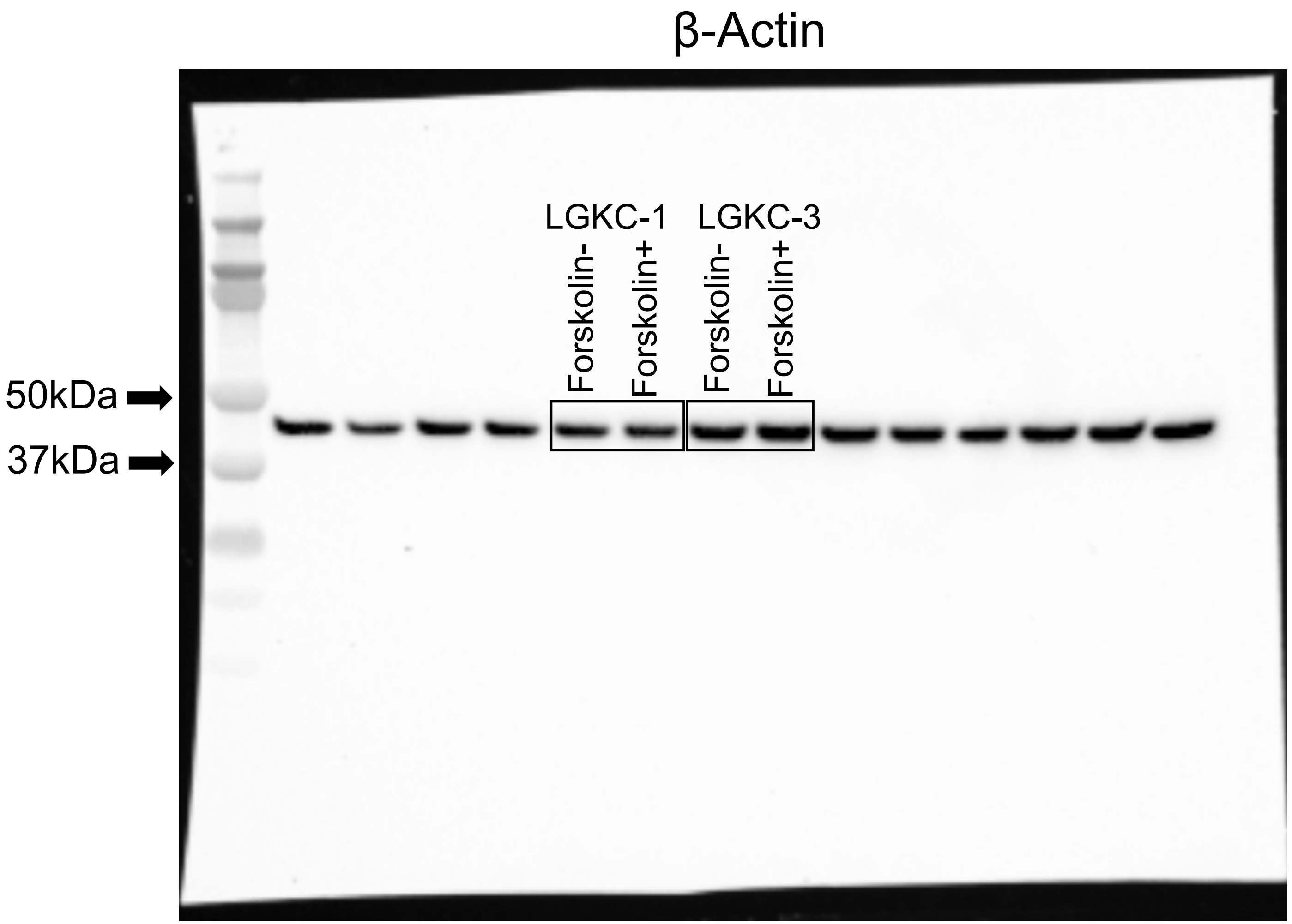
β-Actin
LGKC-1
LGKC-3
Forskolin-
Forskolin+
Forskolin-
Forskolin+
50kDa
37kDa

Figure 7C

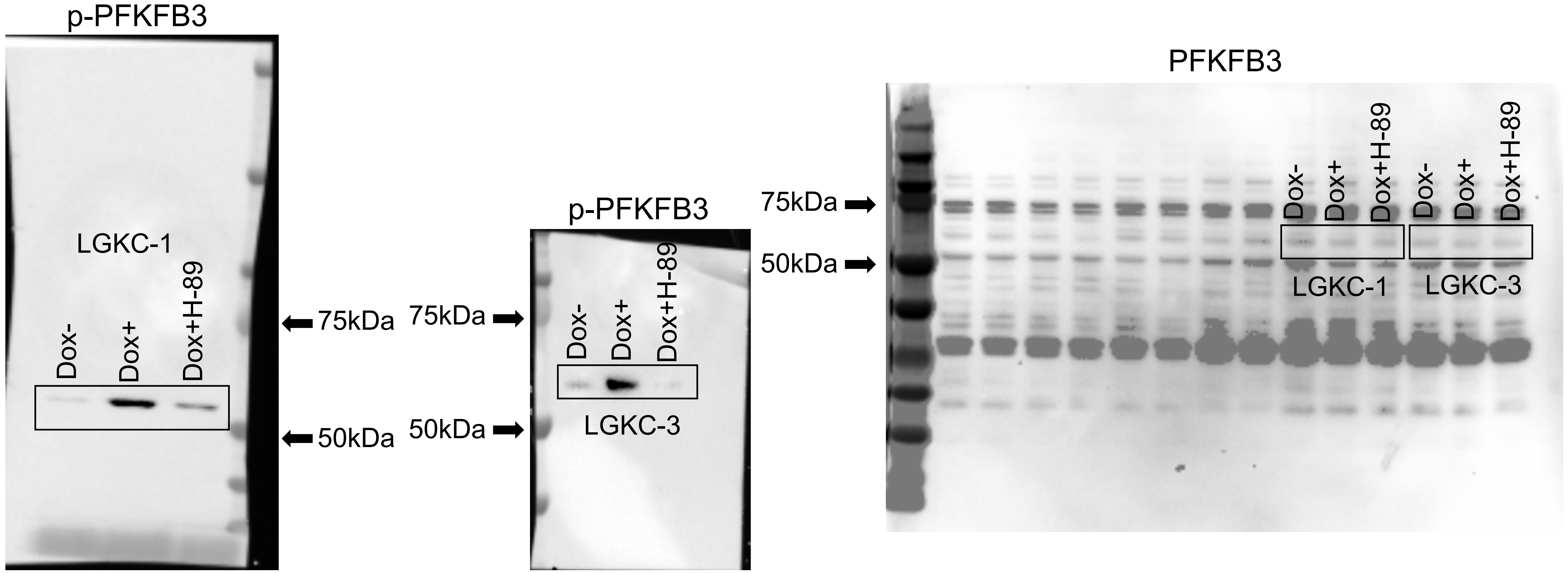
p-PFKFB3
LGKC-1
Dox-
Dox+
Dox+H-89
75kDa
50kDa
p-PFKFB3
75kDa
50kDa
Dox-
Dox+
Dox+H-89
LGKC-3
PFKFB3
75kDa
50kDa
Dox-
Dox+
Dox+H-89
Dox-
Dox+
Dox+H-89
LGKC-1
LGKC-3

Figure 7C

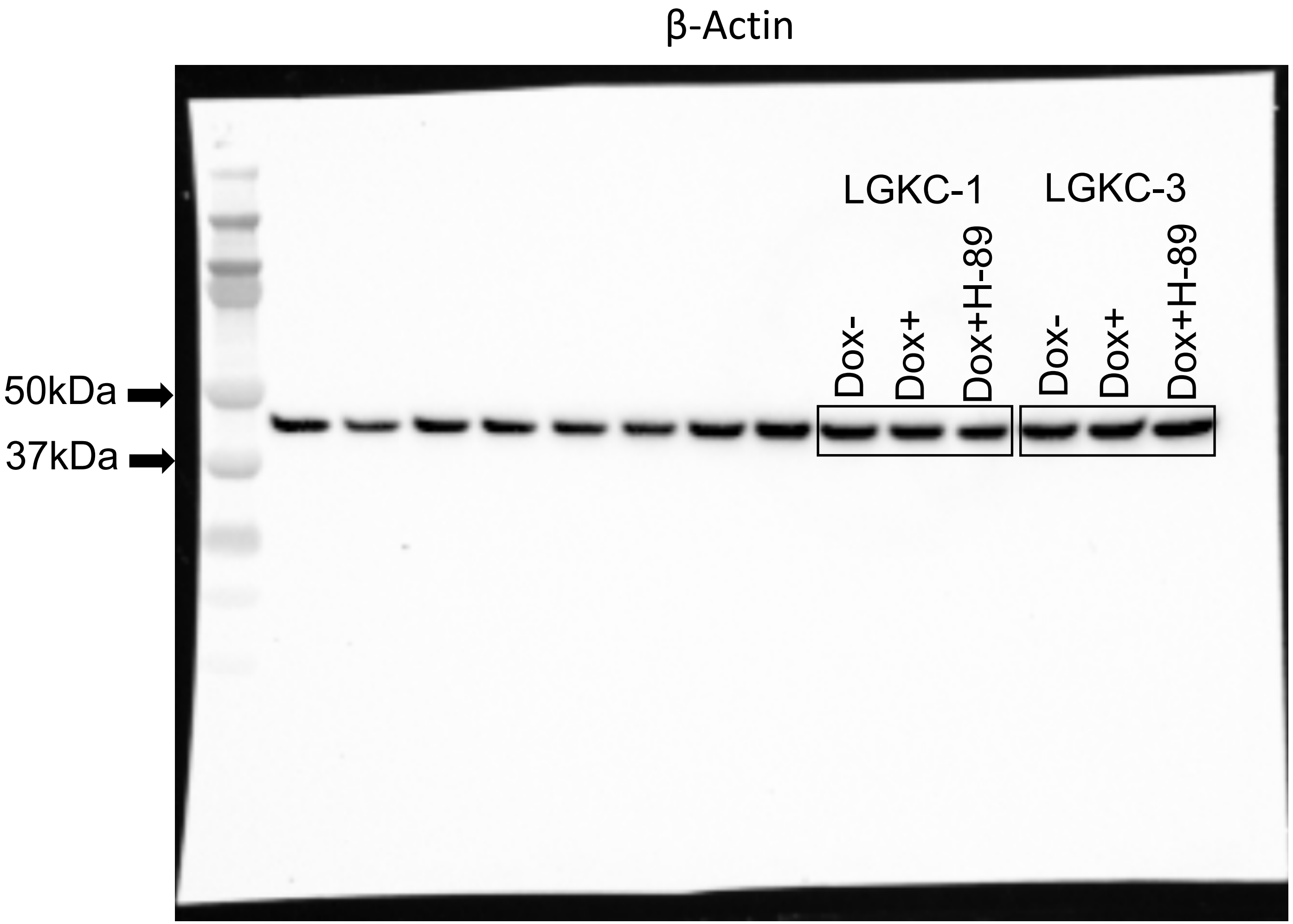
β-Actin
50kDa
37kDa
LGKC-1
LGKC-3
Dox-
Dox+
Dox+H-89
Dox-
Dox+
Dox+H-89

Supplementary Figure 5A

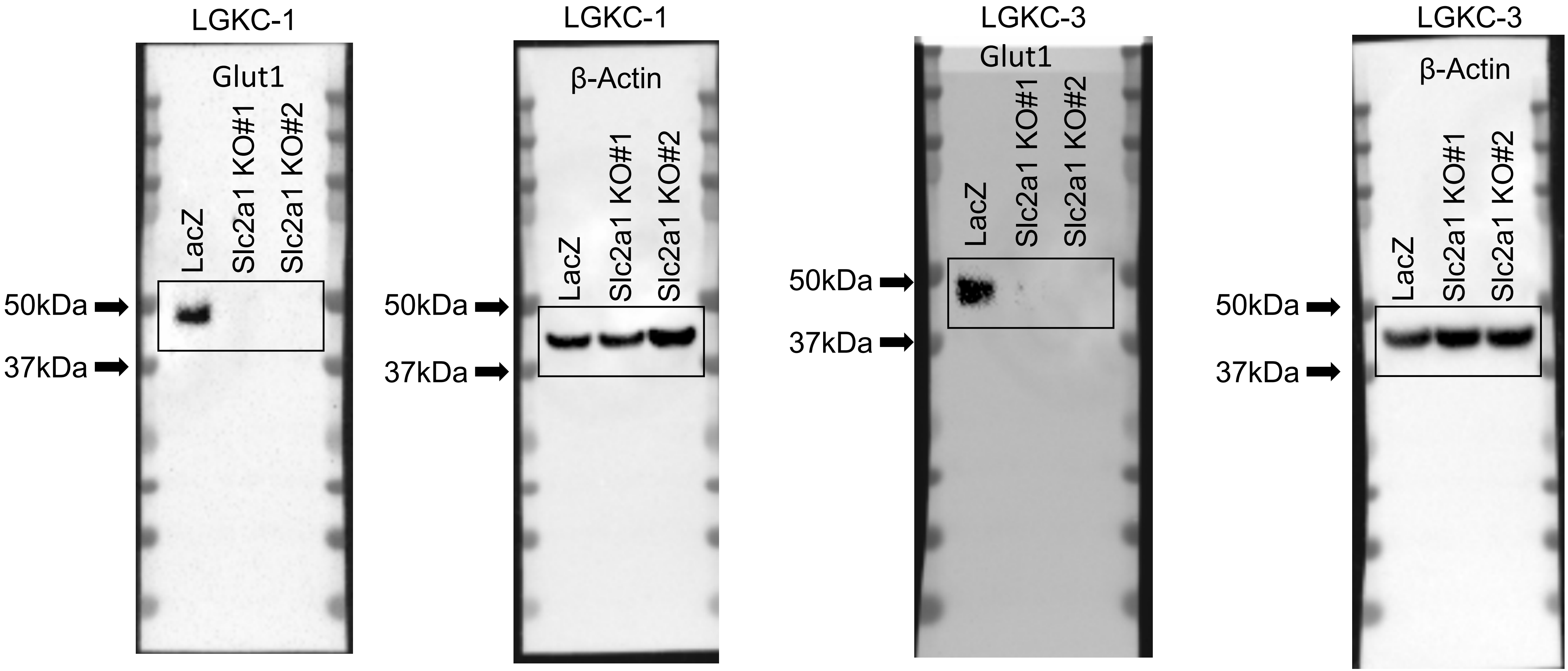
